# Peptide-based inhibitors of Tau aggregation as a potential therapeutic for Alzheimer’s disease and other Tauopathies

**DOI:** 10.1101/2021.06.04.447069

**Authors:** Anthony Aggidis, Shreyasi Chatterjee, David Townsend, Nigel J. Fullwood, Eva Ruiz Ortega, Airi Tarutani, Masato Hasegawa, Hannah Lucas, Amritpal Mudher, David Allsop

**Affiliations:** Division of Biomedical and Life Sciences, University of Lancaster, Lancaster LA1 4YQ, UK; Department of Chemistry, University of Lancaster, Lancaster University, LA1 4YB, UK; Department of Dementia and Higher Brain Function, Tokyo Metropolitan Institute of Medical Science, Tokyo 156-8506, Japan; Department of Biological Sciences, University of Southampton, Southampton SO17 1BJ, UK

## Abstract

There are currently no disease altering drugs available for Tauopathies such as Alzheimer’s disease, which alone is predicted to affect ~88 million people worldwide by 2050. As Tau aggregation underpins its toxicity, aggregation inhibitors are likely to have disease-modifying potential. Guided by in-silico mutagenesis studies, we developed a potent retro-inverso peptide inhibitor of Tau aggregation, RI-AG03 [Ac-rrrrrrrrGpkyk(ac)iqvGr-NH_2_], based on the ^306^VQIVYK^311^ hotspot. Aggregation of recombinant Tau was reduced by >90% with equimolar RI-AG03 and no fibrils were observed by EM. When added during the growth phase, RI-AG03 blocked seeded aggregation. Fluorescein-tagged RI-AG03 efficiently penetrated HEK-293 cells over 24 hours and was non-toxic at doses up to 30 μM. In transgenic *Drosophila*, RI-AG03 significantly improves neurodegenerative and behavioural phenotypes caused by expression of human Tau. Collectively this shows that RI-AG03 can effectively reduce Tau aggregation *in vitro* and block aggregation-dependent phenotypes *in vivo*, raising possibilities for exploring its translational potential.

## INTRODUCTION

There are currently ~54 million people with dementia worldwide, and this figure is predicted to increase to ~135 million by 2050 [1] [2]. Alzheimer’s disease (AD) accounts for 60-70% of these cases. The global societal cost of dementia was estimated at US $1 trillion in 2018 and is predicted to increase to US $2 trillion by 2030 [3]. The average annual cost of managing a person with dementia in the UK is reported as £32,250 [4]. Despite this huge economic burden, the only approved drugs for AD act by modulating neurotransmission to provide temporary symptomatic relief, as opposed to substantially altering the disease course. These drugs include three acetylcholinesterase inhibitors, and an N-methyl-D-aspartate (NMDA) receptor antagonist. This highlights the need to develop improved disease-modifying therapies.

AD is regarded as a proteopathy, manifested through the production of β-amyloid (Aβ) and hyper-phosphorylated Tau proteins that misfold and adopt a high β-sheet content. This initiates a nucleation-dependent aggregation process to form semi-soluble oligomers and eventually fibrils. These fibrillar structures accumulate in the brain as extracellular amyloid plaques, containing Aβ, and intraneuronal neurofibrillary tangles (NFTs), composed of Tau [5] [6] [7] [8]. These protein aggregates are damaging to cells, with the smaller oligomers often being regarded as the most neurotoxic forms of both Aβ and Tau [9]. In the case of Tau, protein monomers stack in register and parallel to each other to form smaller soluble oligomers and ultimately larger insoluble protofilaments which associate in an anti-parallel conformation to form protofibrils and eventually paired helical filaments (PHFs) or straight filaments found in NFTs and other Tau inclusions [10] [11] [12] [13] [14].

There are six isoforms of Tau, dependent on alternative mRNA splicing from the *MAPT* gene on human chromosome 17. The absence or presence of exon 10 determines if Tau has 3 or 4 microtubule-binding repeats. This exon encodes the second repeat, which includes the aggregation hotspot ^275^VQIINK^280^. The other main hexapeptide sequence involved in promoting aggregation is ^306^VQIVYK^311^, found in the third repeat. These hexapeptides are essential for aggregation [15] [16].

The ‘amyloid cascade’ hypothesis proposes that Aβ deposition precedes and initiates NFT formation and has been the mainstream model for AD aetiology for the past two decades [7] and a wide range of data support the notion that Aβ pathology is ‘up-stream’ of Tau pathology [17] [18] [19]. To date, all drugs targeting Aβ have failed stage 3 clinical trials [20] and this has placed more emphasis on Tau. NFT formation in the brain correlates more closely with the destruction of neurones and deficits in memory function than accumulation of Aβ [21]. There is also a move towards combination therapies and a dual-targeting treatment against Aβ and Tau may be more effective than monotherapies [22]. The current compounds investigated in clinical trials for Tauopathies are ACI 35, LMTM, AADvac 1, Davunetide, BIIB076, BIIB080, BIIB092, C2N 8E12, JNJ-63733657, LY3303560, RO7105705, Abeotaxane, UCB0107, CCT020312, Nilotinib, AZP2006, MK-8719, Salsalate, Saracatinib, ASN120290 and IVIg [23] [24] [25]. Only LMTM (TauRx Therapeutics) is listed as a protein aggregation inhibitor, and only NAPVSIPQ (NAP) (Allon Therapeutics/Paladin Labs) as a peptide. There are currently no peptide aggregation inhibitors for Tau in clinical trials. Our research group has previously developed aggregation inhibitor peptides which target aggregation “hotspots” in Aβ and amylin [26] [27]. Using a similar rationale we designed peptide inhibitors to target and mimic aggregation ‘hotspot’ ^306^VQIVYK^311^ in the tau protein sequence, while avoiding peptide self-association and stimulation of Tau aggregation via carefully chosen modifications [28].

## RESULTS

In-silico investigations confirmed suitability of targets in Tau through an informed guidance approach. Aggrescan and Camsol intrinsic algorithms highlighted ^306^VQIVYK^311^ as the region in Tau having the lowest intrinsic solubility, whilst also being prone to aggregation (**Figures S1** and **S2)**.

By conducting residue optimisation to the hexapeptide, aggregation propensity can be reduced to avoid peptide self-association whilst maintaining target specificity. **Table S3** highlights residue optimisations *in silico* conducted on VQIxYK, and VQxVYK, respectively, where ‘x’ was systematically replaced with all possible amino acid residues. Based on the Na4vSS value, Lysine was highlighted as the optimum replacement for both VQxKYK and VQIxYK. It reduced the aggregation propensity to just below the aggregation threshold, suggesting that target affinity would be retained, (**Figure S8)**. His replacement was rejected as its intrinsic insolubility matched that of the native peptide. VQI**K**YK retained some intrinsic insolubility whereas VQ**K**VYK did not, (**Figure S7)**, so was preferred. The substituted Lys was acetylated to render it a hydrogen bond donor to enhance its binding to ^306^VQIVYK^311^ which is in a hydrogen bond acceptor rich region [29] (**Figure S8**).

Experimental peptides were computationally docked to Tau to see if they bound to their target region. **Figure 1** summarises the binding sites and raw data for the average (n=3) of peptide docking to Tau_306-378_. Based on the ICM score, the control peptide [VQIVYK] bound more strongly to the top of the protofilament than modified [VQIK(Ac)YK] did, however with the additional proline, taken from the native sequence, [VQIK(Ac)YKP] bound the strongest. When binding to the bottom of the protofilament however, [VQIK(Ac)YK] bound the strongest.

**Figure 1:**
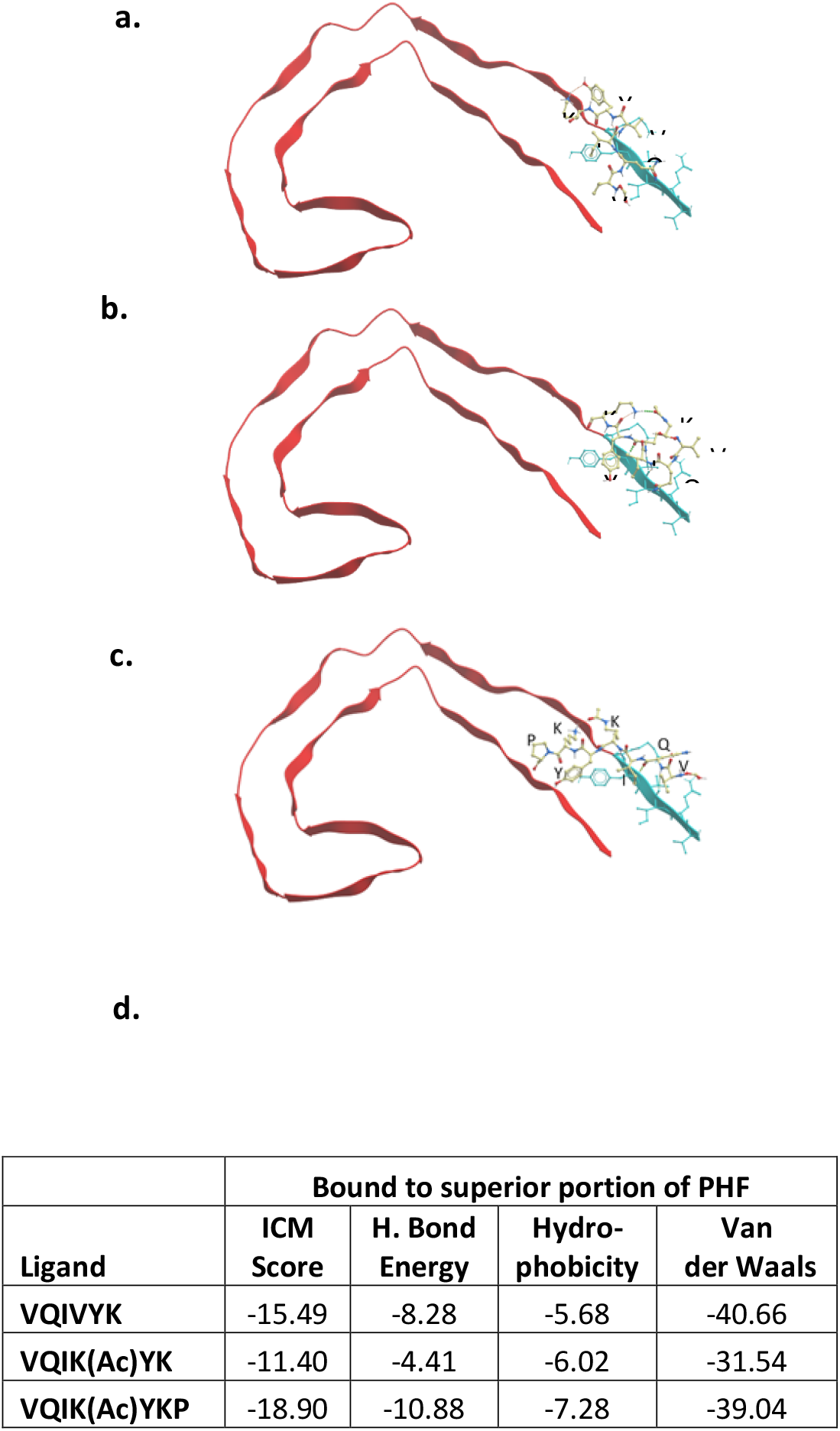
Experimental peptides docked in parallel to the superior portion of PDB 5o3l PHF, demonstrating preferential predicted binding to their complementary ^306^VQIVYK^311^ target (cyan). **a:** VQIVYK; **b:** VQIK(Ac)YK; **c:** VQIK(Ac)YKP, notice the peptide extending to interact with the parallel β-sheet to ^306^VQIVYK^311^; **d**: Summary table of the average computationally calculated values describing the docked compounds to Tau_306-378_. Lower scores indicate more powerful the interactions. **ICM score** of < −32 indicates a very strong bind.

**Figure 2** demonstrates that [VQIVYK] binds in parallel to its complementary sequence in Tau and that it also binds to VQIINK. However [VQIK(Ac)YKP] bound in anti-parallel to its complementary sequence and bound in parallel to VQIINK (also seen in **Figure S9–S10)**. **Table S4** shows that [VQIVYK] and [VQIK(Ac)YKP] both docked to multiple Tau structures from the PDB with similar intensity. The most noteworthy difference in energy was that [VQIK(Ac)YKP] bound almost twice as strongly to the Fitzpatrick PHF structure [10] than native [VQIVYK] did. For referenc e, both heparin and ThT bind to Tau very strongly with ICM-scores of −34.73 and −28.45. A range of peptides (**Table S5**) were designed and tested for their ability to selfaggregate and to inhibit heparin-induced aggregation of TauΔ1-250 *in vitro* (**Figure 3).** These peptides have Ac-RG….GR-NH_2_ flanking sequences to aid solubility, a strategy employed in our previous work [26] [27] [30] [31].

**Figure 2:**
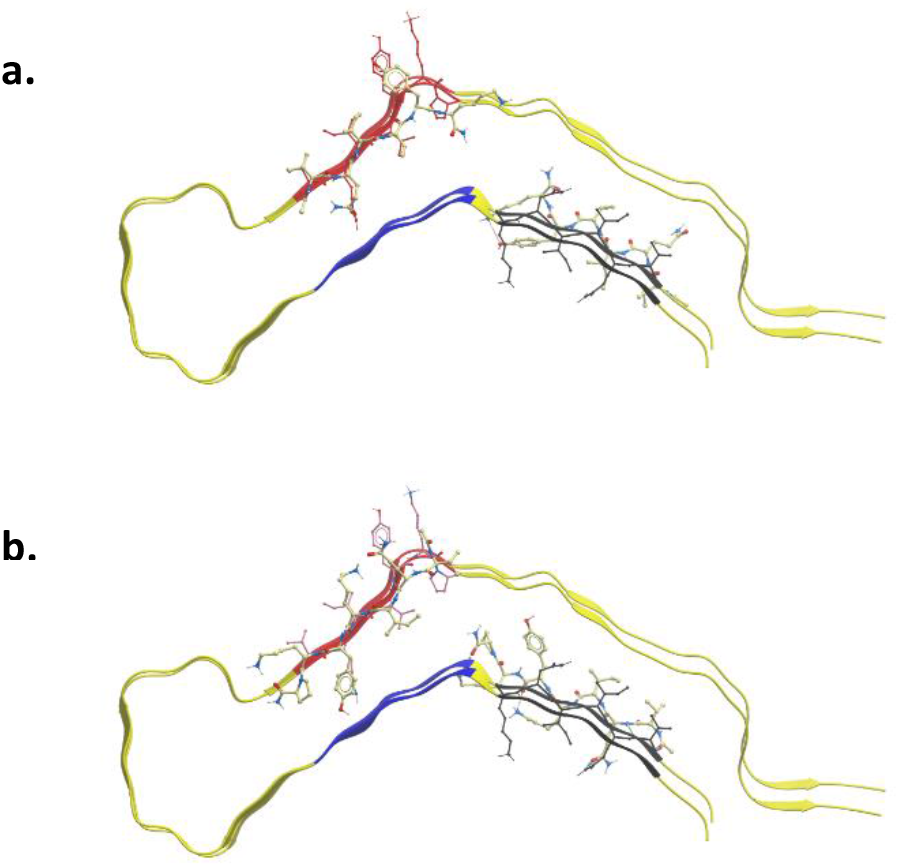
Experimental peptides docked to PDB 6QJH heparin-induced 2N4R Tau snake filament have two predicted binding sites; ^306^VQIVYK^311^ and ^275^VQIINK^280^. **a:** Ac-VQIVYK-NH^2^ binding in parallel to the filament, **b:** Ac-VQIK(Ac)YKP-NH^2^ binding in anti-parallel to the filament at ^306^VQIVYK^311^ position and in parallel at the ^275^VQIINK^280^ position.

**Figure 3:**
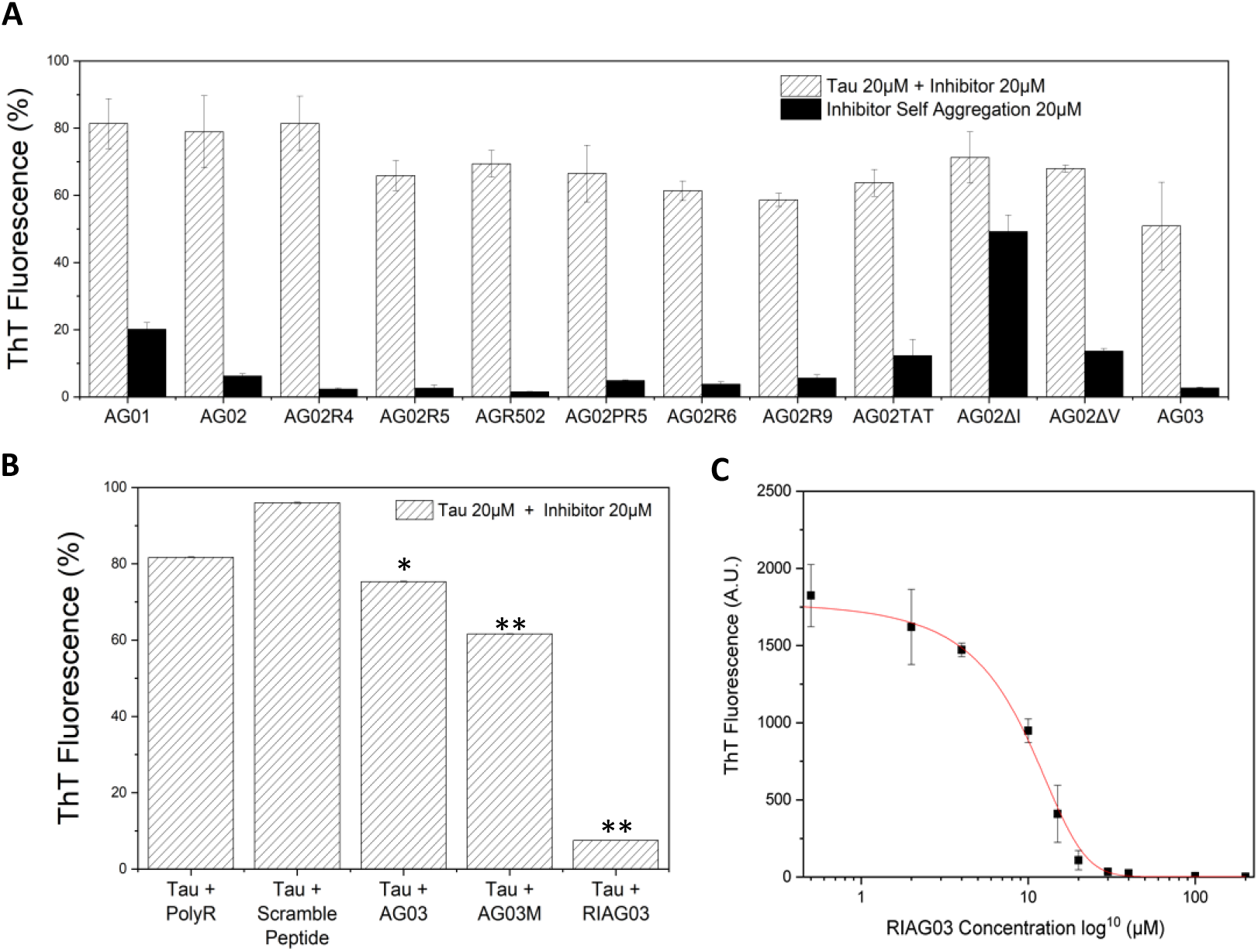
Synergy2 End-point (24 hr) Thioflavin data aggregation of TauΔ1-250 20 μM. **a:** With different peptide inhibitors (white, hatched) and attempted self-assembly of inhibitors without the presence of Tau (black). Key: AG01 [RG-VQIINK-GR], AG02 [RG-VQIVYK-GR], AG02R4 [RRG-VQIVYKG-RR], AG02R5 [RG-VQIVYK-GRRRR], AGR502 [RRRRG-VQIVYK-GR], AG02PR5 [RG-VQIVYKP-GRRRR], AG02R9 [RG-VQIVYK-GRRRRRRRR], AG02TAT [RG-VQIVYK-GRYGRKKRRQRRR], AG02ΔI [RG-VQK(Ac)VYK-GR], AG02ΔV [RG-VQIK(Ac)YK-GR], AG03 [RG-VQIK(Ac)YKP-GRRRRRRRR]. **b:** With equimolar concentrations of either octa-arginine, scrambled AG03 peptide, AG03, N-methylated AG03 or retro-inverso AG03. n=3, error bars reported as standard deviation. **c:** Dose dependent log10 scatter graph employing a curve fitting algorithm to calculate the IC50 of 7.83 μM. * P ≤ 0.05, ** P ≤ 0.01, *** P ≤ 0.001.

Peptides AG01 [Ac-RG-VQIINK-GR-NH_2_] and AG02 [Ac-RG-VQIVYK-GR-NH2] self-aggregated, and so were deemed to be poor choices as inhibitors (**Figure S11**). VQIVYK based peptides were focused on for development as this motif is present in all Tau isoforms. **Figure 3** demonstrates that increasing solubility via the addition of up to three consecutive Arginine residues inhibited aggregation by an additional 20%, presumably through charge effects and steric hindrance. In addition, the inclusion of a total of 9 Arginine residues (AG02R9) reduced aggregation further by an additional 10%. As Tau aggregates intracellularly it is important that the inhibitor has a cell penetrating sequence so R9 and TAT were tested as these are required to aid cell and brain penetration [32]. Inclusion of the TAT sequence (AG02TAT) did not result in any additional inhibitory benefits compared to AG02R5 and AG02R9 self-associated less than AG02TAT. The termini these poly-Arginine chains were located on did not greatly change peptide inhibitory action. No additional benefit was observed when introducing Proline to the end of the binding sequence (AG02PR5), however it was retained due to it being found in the native VQIVYKP sequence. The hypothesis was that angle change in concert with a longer poly-Arginine chain would create additional interference for other Tau molecules. AG02ΔI [Ac-RG-VQK(Ac)VYK-GR-NH_2_] demonstrated self-seeding ability which agreed with the docking predictions. AG02ΔV [Ac-RG-VQIK(Ac)YK-GR-NH_2_] suggested a tendency for self-aggregation. However, AG03 [Ac-RG-VQIK(Ac)YKP-GRRRRRRRR] this inhibited aggregation the most (~53% inhibition), while showing minimal self-aggregation. Thus, AG03 was selected as the ‘lead’ inhibitor for further development. The difference between the binding sites of AG02 and AG03 include the replacement of the second Valine with Acetyllysine and adding Proline on the end for AG03.

AG03 needed to be proteolytically stable so an N-methylated (AG03M) [Ac-RG-V(m)QI(m)K(Ac)Y(m)KP(m)-GRRRRRRRR-NH_2_] and a retro-inverted version (RI-AG03) [RI-RG03 = Ac-rrrrrrrG-pkyk(ac)iqv-Gr-NH_2_] were tested against Tau aggregation. AG03M showed no improvement over its parent molecule, but RI-AG03 inhibited Tau aggregation by ~94% at equimolar concentration with an IC50 of 7.83 (**Figure 3c**). Octa-Arginine [RRRRRRRR] and scrambled AG03 [Ac-RG-QPKIK(Ac)YV-GRRRRRRRR] controls which do not include the binding region had no effect, demonstrating that inhibition was not solely due to charge effects and emphasizing that it is not the presence of the amino acid residues, but also their sequence, which endows the inhibitory properties. **Figure 3c** demonstrates the dose-dependent relationship of RI-AG03 against 20 μM TauΔ250 aggregation, with IC50 = 7.83 μM.

RI-AG03 demonstrated efficacy at inhibiting seeded aggregation when added one hour into the growth phase of aggregation and halting any further aggregation (**Figure 4a**).This confirmed that previous inhibition of Tau aggregation was not caused by the inhibitor’s positive charges sequestering heparin. The aggregation of Tau alone and in the presence of the inhibitor RI-AG03 was studied using time resolved circular dichroism. Tau filament formation is expected to coincide with a reduction in disordered structure and an increase in β-sheet content [33]. The specific construct used in this study has previously been shown to follow this transition with heparin induced aggregation [34] [35]. Secondary structure estimation from the CD spectra using the Bestsel algorithm [36] [37] reveals this Tau construct has a predominantly disordered native structure (56 %), with the rest made up of turns (18 %), and β-sheet (26 %) (**Figure 4B red**). Upon incubation under aggregation inducing conditions, the unstructured content and turns decreases over 5 hours to 48 % and 15 %, respectively, with an increase in β-sheet to 31 % and α-helical content to 6 %. No further change occurs between 5 and 24 hours. Incubation of Tau with RI-AG03 results in a similar native spectra and estimated structure (disordered 57 %, turns 18 %, β-sheet 23 % and α-helical 2%) (**Figure 4B blue**). This remains largely unchanged after 5 hours, (disordered 56 %, turns 17 %, β-sheet 22 %, and α-helical 5 %) with crucially, no increase in β-sheet structures as seen for the untreated tau. This suggests that RI-AG03 impairs Tau self-assembly by stabilising the Tau in a structure close to its native form. When TauΔ1-250 was incubated with heparin for 24 hours, insoluble fibrils typical of amyloid were distributed evenly across the TEM surface (length and diameter). However, in the presence of equimolar RI-AG03 these fibrils were not observed, and only very small spherical structures, with mean diameter ~36 nm, were present (**Figure 4e**).

**Figure 4:**
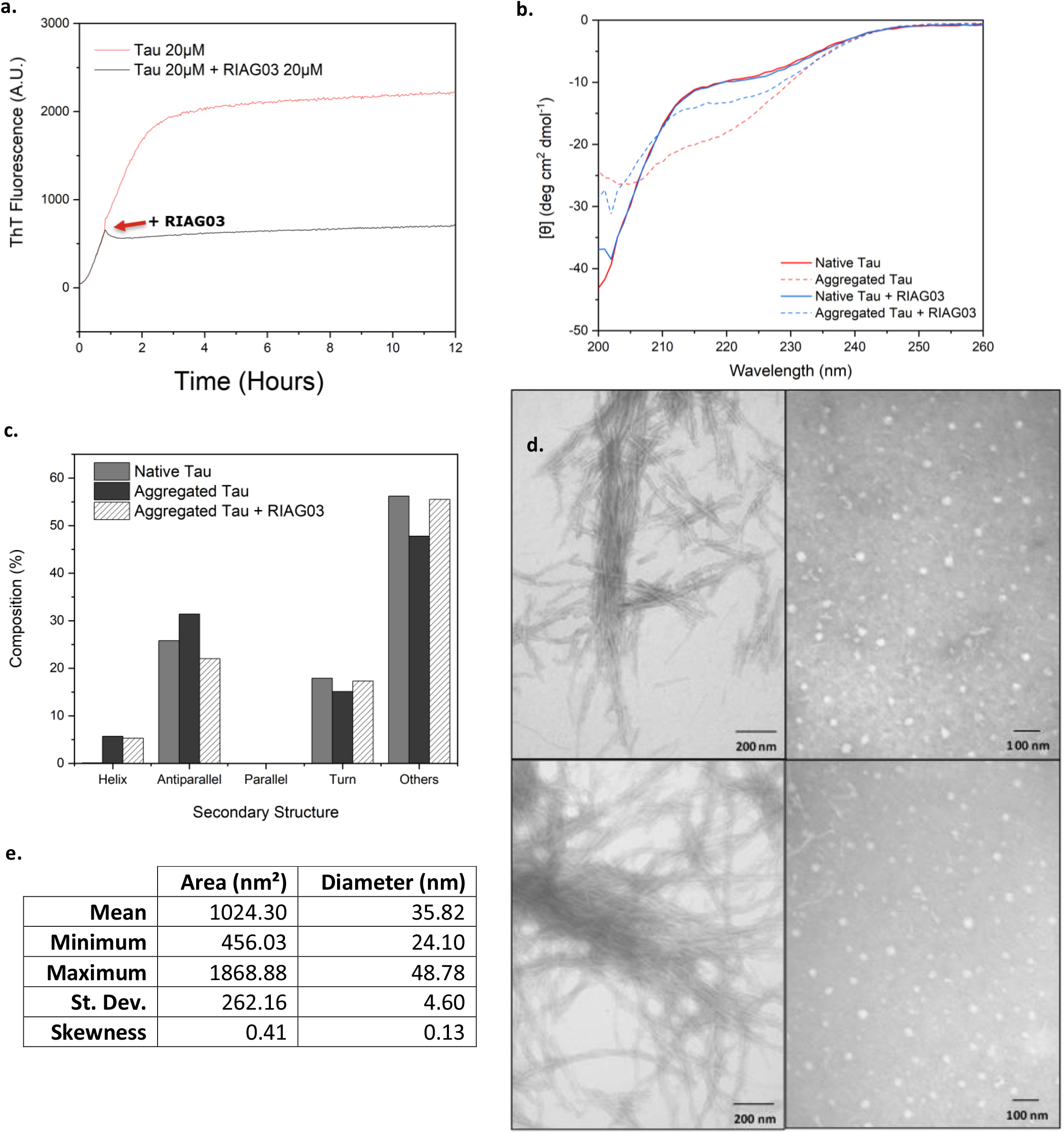
ThT Kinetics, CD spectroscopy and EM data **a:** Aggregation kinetics of TauΔ1-250 20 μM with RI-AG03 20μM added one hour into the experiment. **b:** Incubation with heparin and RI-AG03 at 37°C for 5 hours; TauΔ1-250 20μM demonstrates a reduction in β-sheet content (at 220 nm) compared to control; **c:** Secondary structure estimation analysis of 4b using BeStSel; **d**: Negative stain TEM images using a JEOL JEM-1010 following aggregation of TauΔ1-250 20 μM at pH 7.4 in the presence of Tris buffer 30 mM, DTT 1 mM, heparin 5 μM incubated at 37°C for 24 hr, without (left) and with RI-AG03 20 μM (right). Note absence of fibrils in the presence of RI-AG03. Repeats were performed in triplicate across independent experimental repeats; **e:** Table quantifying area and diameter of 200 oligomeric-like structures seen in TEM images from Tau incubated with RI-AG03, using iTEM software.

Unlike AG03, RI-AG03 showed resistance to digestion with Trypsin over 24 hours, as shown by SDS-PAGE (**Figure 5**). RI-AG03 penetrated HEK-293 cells and was non-toxic even at doses up to 30 μM over 24 hours (**Figures 5a and b)**. This provides a degree of confidence for the potential translational utility of this peptide prior to experiments *in vivo* in a *Drosophila* model of tauopathy. BLASTP 2.8.1+ shows no match for human Tau residues ^275^VQIINK^280^ and ^306^VQIVYK^311^ in *Drosophila* Tau, suggesting that RI-AG03 will not interfere with endogenous Tau in this model.

**Figure 5:**
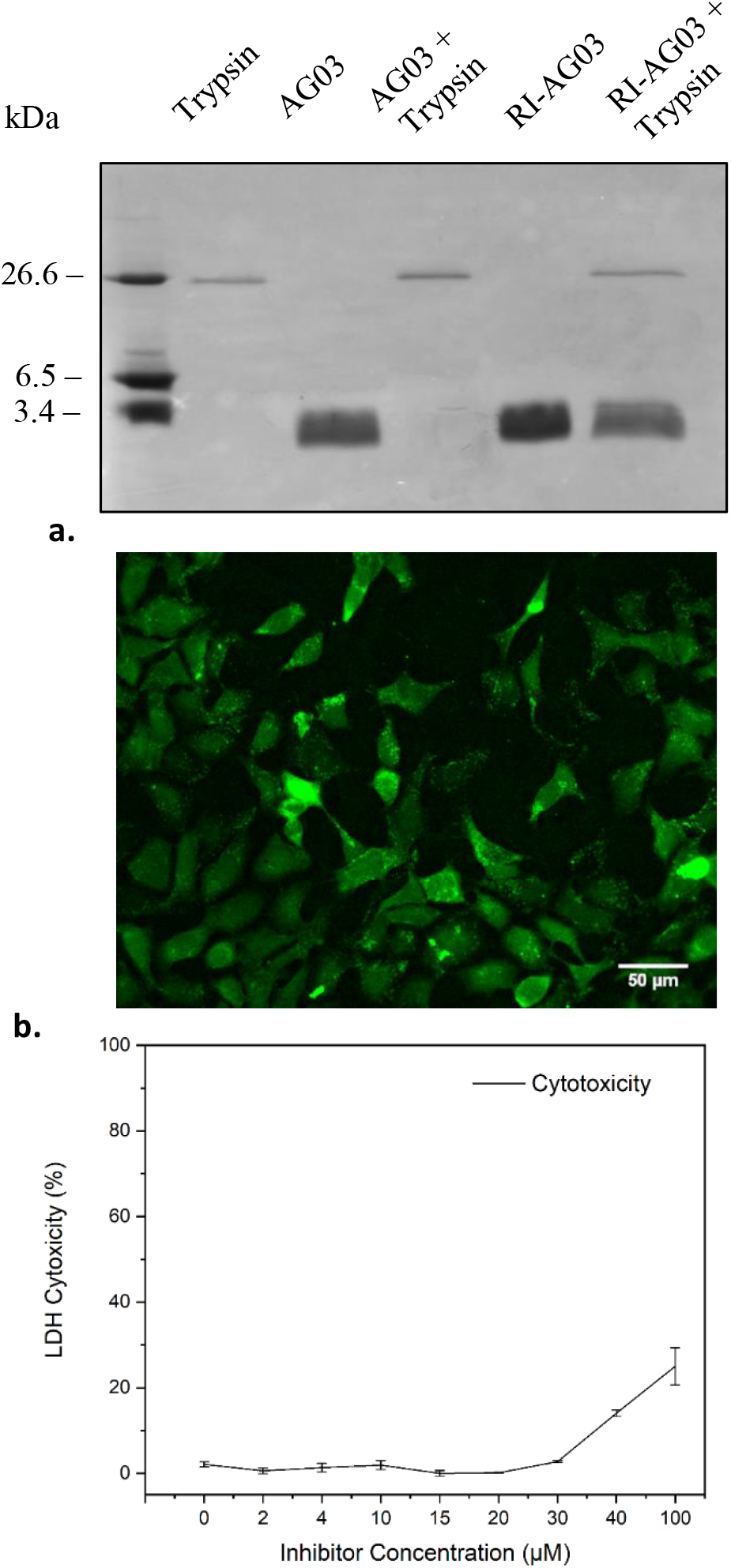
**A:** 15% SDS PAGE gels of AG03 (100 μM) and RI-AG03 (100 μM) treated with and without equimolar Trypsin concentration for 24-hours at 37°C. AG03 has no signal in the presence of Trypsin whereas RI-AG03 has a strong signal in the presence of Trypsin. Densitometric analysis suggested a 19% reduction in signal after equimolar incubation of RI-AG03 with Trypsin. **B:** Cellular uptake of FAM-RI-AG03 (15μM) by HEK-293 cells over 24-hours. Almost no peptide is visible in the medium, suggesting that majority of the peptide was taken up into the cells. **C:** Varying concentrations of RI-AG03 were co-incubated with HEK-293 cells and cytotoxicity was analysed using an LDH cytotoxicity kit. Toxicity begins to increase at 40μM.

RI-AG03 was fed to an established *Drosophila* model of Tauopathy overexpressing full-length human Tau (2N4R) in the eye using the eye-specific *GMR*-Gal4 driver. These human-Tau overexpressing flies undergo profound Tau-induced neurodegeneration in the photoreceptors which is reflected in a reduced eye-size compared to controls (**Figure 6**). Eye-size was restored in Tau-transgenics when treated with 40 μM RI-AG03, showing a statistical difference compared to the untreated Tau flies (**Figure 6b**). SEM images of the Tau-transgenics further displayed an array of fused ommatidia and missing bristles in the anterior part of the eye leading to the characteristic Tau-induced “rough-eye” phenotype. RI-AG03 treatment partially rescued this “rough-eye” phenotype by improving the number and morphology of bristles and reducing the number of abnormal ommatidia as seen by SEM (**Figure 7).** Inhibitor**-**treated *Drosophila* appeared more like controls and demonstrated a significant improvement in overall eye-morphology and ommatidial organisation when compared to untreated Tau-transgenics.

**Figure 6:**
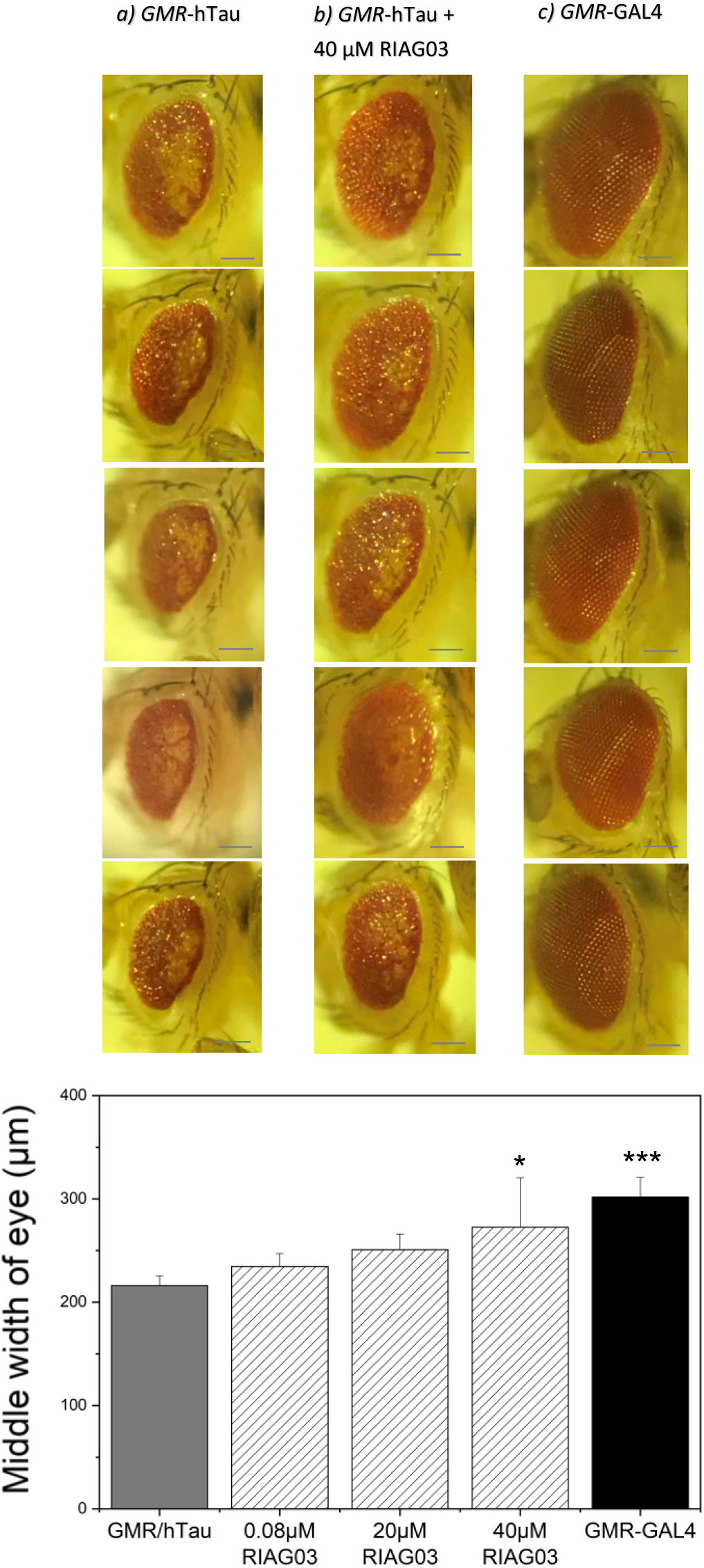
Shows experimental data of *Drosophila* overexpressing Tau in the eye (GMR-hTau) treated with various concentrations of RI-AG03 and healthy *Drosophila* control (GMR-GAL4). Tau expression causes toxicity resulting in morphological changes in the eye width at the middle of the eye. Notice the increased width of the eye when the Tau fly (**a.**) is treated with RI-AG03 (**b.**), closer resembling GMRGAL4 (**c.**). Scale bar = 100 μm. **d:** Width of the eye at the middle increased with 40 μM treatment Data presented as means (n=5/condition) and standard deviation. One factor repeated measures ANOVA + Tukey post hoc statistical analysis was conducted.

**Figure 7:**
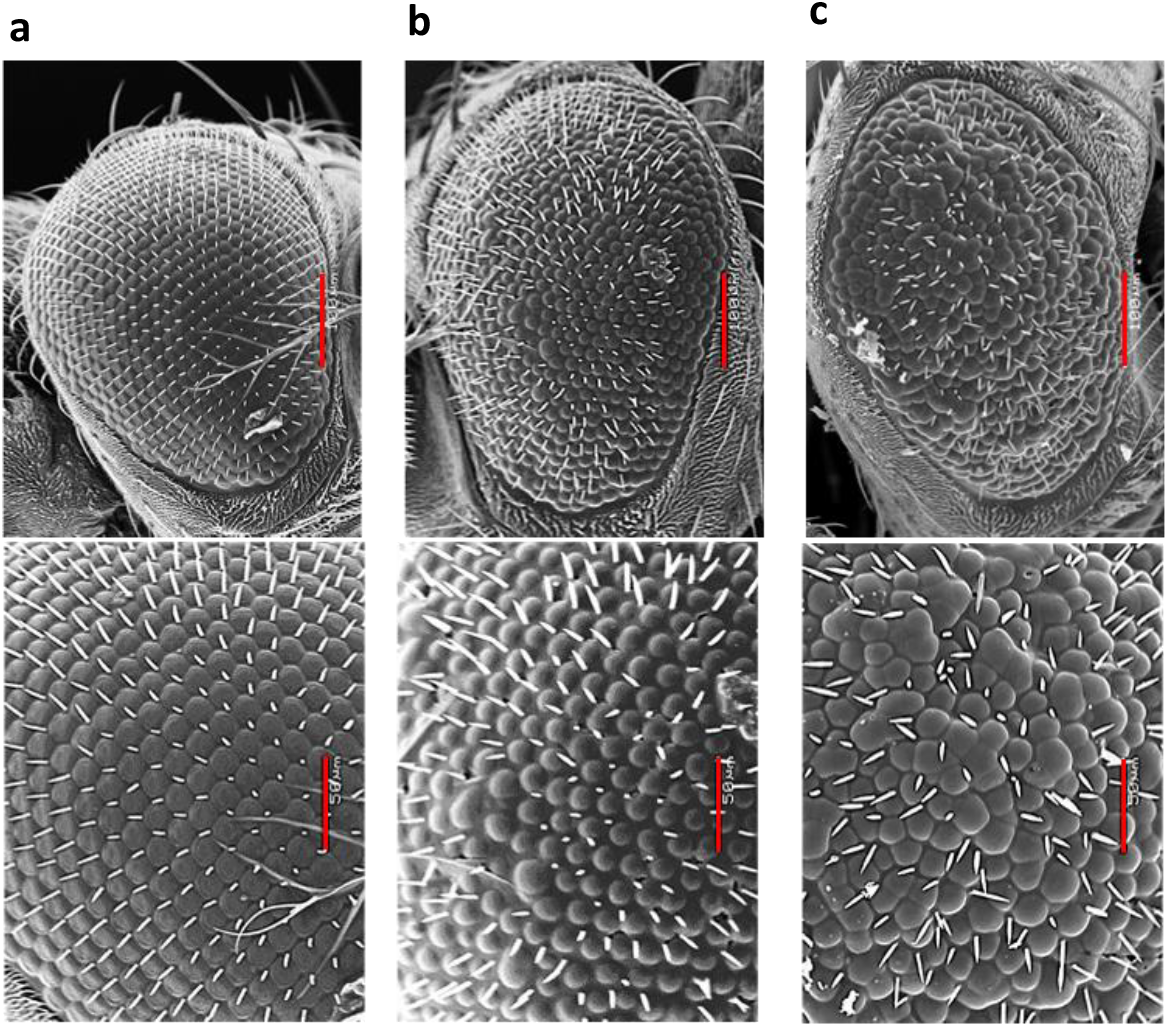
*Drosophila* eye SEM data of (panels left to right) **(a)** healthy *GMR*-GAL4, **(b)** *GMR*-hTau treated with 20μM RI-AG03 and **(c)** *GMR*-hTau without treatment. Upper panel (scale bar = 100 μm): The treatment with 20 μM RI-AG03 restores the shape and size of the Tau-induced rough eye in **(b)** compared to untreated Tau transgenics **(c)**. Lower panel (scale bar = 50 μm): Treatment with 20 μM RI-AG03 restores the number of ommatidia in *GMR*-hTau compared to untreated Tau transgenics where fused ommatidia and missing bristles are visible.

To test whether this rescue of Tau-induced rough-eye phenotype could be extended to ameliorate Tau-induced neuronal dysfunctions in whole organisms, the impact of RI-AG03 on lifespan of flies expressing 2N4R pan-neurally was assessed. Tau-expressing (Elav/Tau) and control (Elav/Orer) flies were reared on media containing either low-dose (0.08 μM) or high-dose (0.8 μM) of the RI-AG03 inhibitor and survival assays were performed [38]. Whilst the lifespan of control flies varied from 80 to 90 days and was unaltered by inhibitor treatment, the lifespan of the Tau flies was significantly improved by treatment with both high and low dose of inhibitor (**Figure 8**). Median survival of Tau flies increased from 26 to 35 days following inhibitor treatment (p< 0.0005 n=100) whilst that of controls was not significantly altered (67 to 70; p>0.05 n=100). This suppression of two Tau-induced phenotypes, known to be Tau aggregation-dependent [39] suggests that RI-AG03 has efficacy as a suppressor of tau-mediated toxicity *in vivo*.

**Figure 8:**
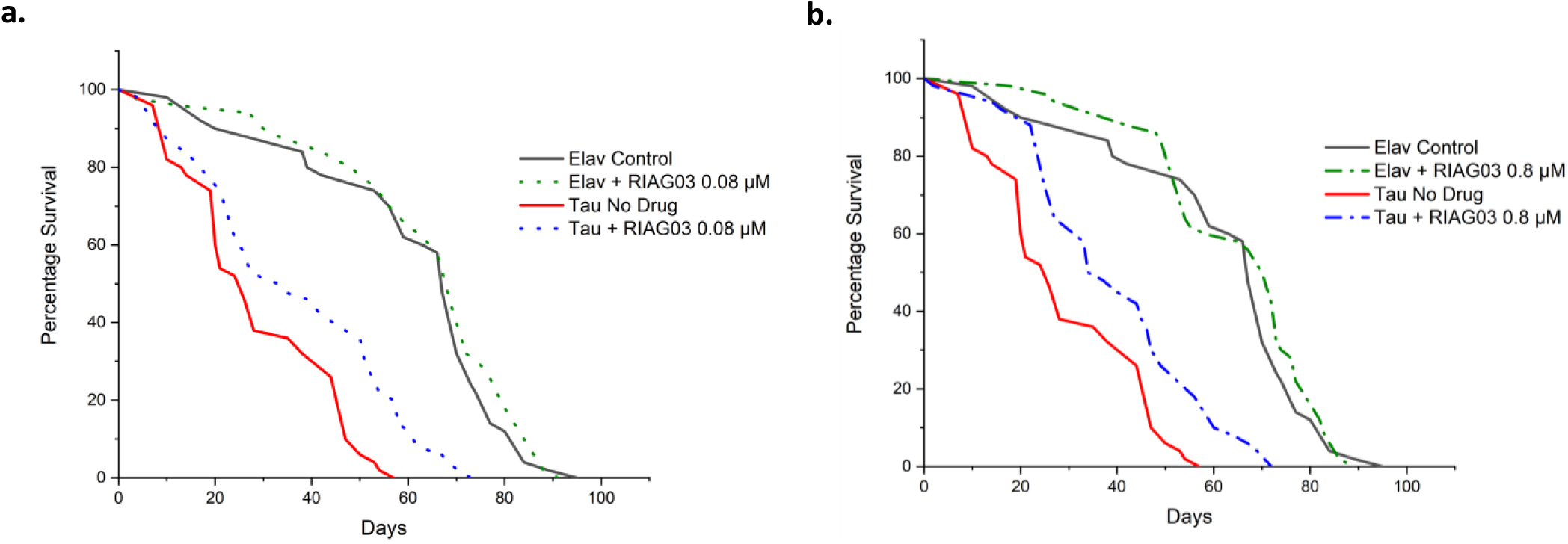
Survival curves of Control (Elav-GAL4) and Tau overexpressing flies (Elav-hTau) in the presence and absence of low (0.08 μM) and high (0.8 μM) doses of the RI-AG03 respectively. **a:** Treatment with 0.08 μM and **b:** 0.8 μM RI-AG03, significantly increased the lifespan of hTau expressing flies by approximately 2 weeks, p = 0.0007 and 0.0004, respectively, while that of treated controls remained unchanged p = 0.194 and 0.332, respectively (Log-Rank test n=100 per genotype/treatment group). Median survival rates were increased from 26 in untreated Tau flies to 33 and 35 after the low and high dose inhibitor treatment, respectively. In Elav controls the median survival rates remained unchanged after the low and high dose inhibitor treatment (67 to 70).

Inhibitory properties of RI-AG03 were improved when covalently attached to liposomes, reducing TauΔ1-250 aggregation ThT fluorescence by ~94% (**Figure 9**). Control samples of liposomes without RI-AG03 had no inhibitory properties.

**Figure 9.**
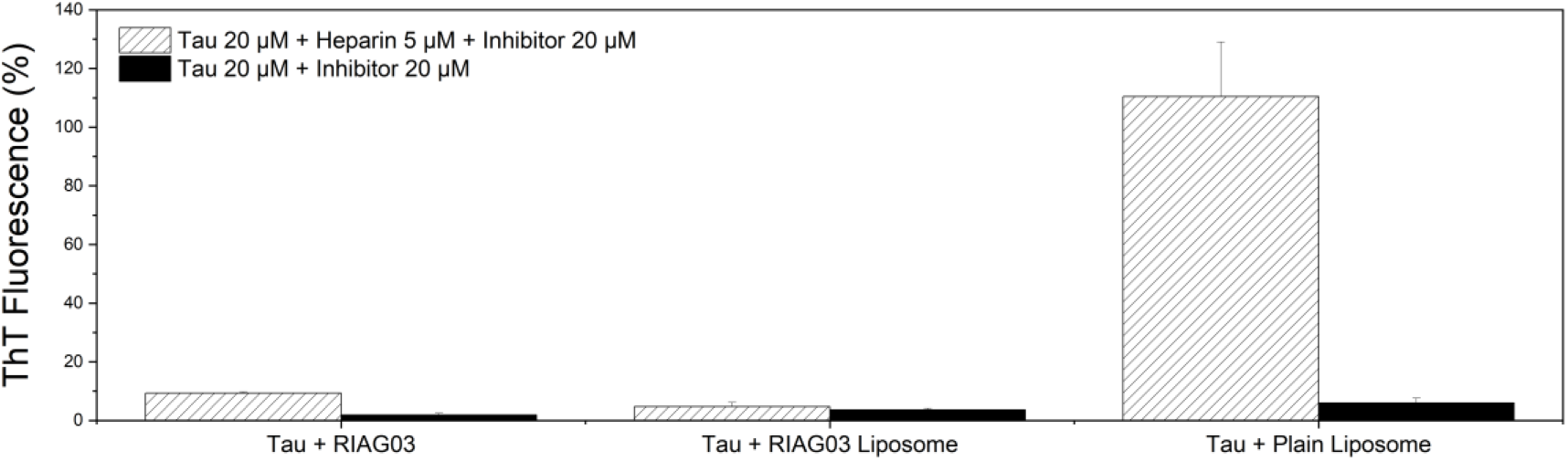
Thioflavin data using FlexStation 3. **a**: Heparin induced aggregation of TauΔ1-250 in the presence of equimolar concentrations of either RI-AG03 or RI-AG03 Liposomes (white) or without (black); **b**: Heparin induced aggregation of TauΔ1-250 in the presence of equimolar concentrations of plain liposomes (white) and without (black). Concentration of liposomes was calculated from the molecular mass of the components of the liposome.

## DISCUSSION

Deletion of ^306^VQIVYK^311^ inhibits pathogenic aggregation of Tau, whereas co-incubation of Tau with this peptide stimulates aggregation [28]. This, in concert with Aggrescan and Camsol results (**Figure S1–2)** emphasises the importance of this sequence in Tau aggregation as a target for therapeutic action. We describe here the development of one such peptide and show that it potently inhibits tau aggregation *in vitro* and significantly ameliorates aggregation-dependent Tau phenotypes *in vivo.*

As VQIVYK stimulates aggregation, this sequence was used for designing the aggregation-inhibiting peptide. First, modifications were required to prevent peptide self-association. Isoleucine^308^ and Valine^309^ were chosen for mutagenesis because they are hydrophobic amino acids within the core of the hexapeptide known to influence aggregation [33]. Tyrosine was not altered as it is useful as an anchoring residue by aromatic stacking interactions. Only one location in the hexapeptide was changed to avoid losing target specificity.

Not all hydrophilic amino acid substitutions were able to reduce the aggregation propensity. There is a fine line between functional and pathogenic protein behaviours, and it is logical to assume that a Na4vSS value close to but below the aggregation threshold would be desirable. The rationale was that if the value was too low, then the inhibitor would yield a weak interaction with its target, but too high and it may seed aggregation. Isoleucine^308^ plays an important role in hydrophobic interactions, supported by its sidechain β-carbon restricting its conformations and involvement in binding recognition [40]. In addition, Valine^306^ and Isoleucine^308^ form an apolar dry interface with Tyrosine^310^ and pack tightly with the 4^th^ repeat [10] [40]. CH/π interactions between Isoleucine and Tyrosine in ^306^VQIVYK^311^ are essential for dry interface formation of steric zippers [41]. These interactions were observed here through molecular docking simulations.

Due to its bulky pyrrolidine ring, Proline is a β-sheet breaker as it places steric conformational constraints on preceding residues with its fixed dihedral angle of −65° which is incompatible with β-sheets [42] [43]. Substitution of Proline into the sequence was not deemed appropriate as its rigidity would interfere with hydrogen bonding between the inhibitor and its target [44]. Instead, Proline^312^ was included in its natural position in Tau. This gives a ‘kink’ in the structure when poly-Arginine is attached to the Proline side of the molecule, which may be beneficial at repelling interacting Tau molecules.

The peptide ^306^VQIVYK^311^ was favoured as it is present in all Tau isoforms and shares similarity to ^275^VQIINK^280^ which might promote binding to both targets [45]. Soft acid/base interactions are important for recognition and assembly, however acetylenic groups form CH/π bonds on par with the strength of classic hydrogen bonds [46]. For Tau molecules to bind and form stable filaments, the positive charges must be neutralised [42]. By removing the positive charge on Lysine, interaction between the inhibitor and Tau is encouraged [47]. The substituted acetylated Lysine also packed tightly with the 2^nd^ repeat which houses the VQIINK sequence [42]. Interference with this region may provide additional disruption to the pathogenic folding of Tau by interfering with how VQIINK folds.

Within the same β-sheet layer VQIVYK organises into parallel face-to-face β-strands. Protofilaments are then formed from mating identical anti-parallel β-sheet layers [48] [49] [50]. In line with this, VQIVYK docked in parallel to its complementary sequence in Tau and the modified peptides [VQIK(Ac)YK] and [VQIK(Ac)YKP], did as well (**Figure 1**).

Free VQIVYK peptides have been shown to form β-sheets of parallel and anti-parallel strands which stack and form disordered/irregular protofilaments [51]. This may explain why VQIK(Ac)YKP docked in an anti-parallel fashion as seen in **Table 4**. Peptides promoting anti-parallel alignments result in slower aggregation kinetics due to increased effort to mate β-sheets [49] [52]. VQIK(Ac)YKP also bound to VQIINK, perhaps due to the peptide sharing the same VQIxxK sequence as these targets. VQIVYK interacts with VQIINK to produce twisted PHF-like filaments [53]. Asparagine^279^ from VQIINK (which can donate 2 hydrogen bonds) may form a quadrupole moment by interacting with Tyrosine^310^ from VQIVYK via polar-π interactions [54]. This anti-parallel interaction may alter the aggregation pathway. When VQIK(Ac)YKP targeted ^305^SVQIVYKP^312^ it bound to residues Glycine^302^, Histidine^374^, Asparagine^286^, Serine^285^, Glutamine^288^, whereas when it targeted ^274^HVQIINKK^281^ it bound to Leucine^282^, Asparagine^297^, Tyrosine^310^, Serine^316^. Interestingly these conformations bind to residues found in both binding sites, specifically Histidine^374^ and Tyrosine^310^ suggesting that the inhibitor may interact with both VQIVYK and VQIINK simultaneously. Using a combination of AG02ΔI and AG02ΔV was not favourable because this would result in a loss of specificity by altering the hydrophobicity and hexapeptide recognition sequence too much. Addition of Arginine residues may enhance inhibition by causing steric hinderance, similarly like N-methylation does.

Another peptide inhibitor developed by our group which outcompetes monomers for the ^16^KVLFF^20^ region in Aβ and named OR-2 [H_2_N-RG-KLVFF-GR-NH_2_], retained its ability to inhibit Aβ aggregation when retro-inverted to RI-OR2 [Ac-rG-ffvlk-Gr-NH_2_], but unlike the comparison between AG03 and RI-AG03, was not a substantially better inhibitor [26] [54]. Furthermore, when the amylin aggregation inhibitor targeted at ^13^ANFLVH^18^ and named IO8 [H_2_N-RG-ANFLVH-GR-NH_2_] was converted to RI-IO8 (Ac-rGhvlfnaGr-NH2), this caused it to actually promote amylin aggregation [27]. N-methylation of IO8 did not have any effect on inhibitory action against amylin aggregation [27], which is what was seen when comparing AG03 and AG03M. It is not predictable whether retro-inversion or N-methylation will necessarily improve the inhibitory effects of these peptide inhibitors. Changes to charge based on the terminal domains and the backbone of peptides can either promote seeding or improve aggregation inhibition. L and D-amino acids are optical isomers, however when peptide sequences are retro-inverted, although their side chains are in similar orientation, their backbones are reversed. This means that the hydrogen bonds formed are different. Amyloid fibre stability is maintained through backbone interactions, so it is striking that RI-AG03 with a reverse backbone has greatly improved activity. Also, terminal modifications can generate new interaction surfaces and enhance stability and activity. The amide functional group when near valine may enhance inhibitory properties by affecting local hydrophobicity [55]. Capping the termini can enhance aromatic interactions, which is useful for the tyrosine residue in the inhibitor [56].

TauΔ250 control aggregated to from ordered structures consisting of PHFs and straight fibrils. However, in the presence of RI-AG03 no fibrils were formed and instead, ~36nm diameter spherical structures were seen. This size is compatible with formation of large granular Tau oligomers (GTO) [57]. It was suggested in a *Drosophila* model that GTO’s are not toxic and may be neuroprotective [58] [59]. Reduction of ThT signal suggests a reduction of Tau β-sheet content, but this method cannot detect the presence of oligomers. Through secondary structure estimation analysis, RI-AG03 normalised β-sheet content to similar levels seen in native Tau, as shown by CD spectroscopy [60] [61] [62] [63].

Heparin induced 2N4R filaments adopt different conformations including twister, snake, jagged and hose filaments which all have a subtle 20° difference to the backbone of the ^310^Tyrosine-Lysine-Proline^312^ residues, resulting in different conformations of β-sheet packing. It is possible that RI-AG03 interferes with this specific region in Tau, but the inhibitor was able to stop Tau aggregation in the absence of heparin. Whether the GTO-like structures are formed when the inhibitor blocks ‘seeded’ aggregation of Tau, in the absence of heparin, remains to be seen.

When RI-AG03 was attached to liposomes it was ~ 40 times more effective as an inhibitor because the concentration of liposome drug was based on the total lipid (not peptide) content. A 1:1 ratio of tau to total lipid was used. RI-AG03-C binds to DSPE-PEG-MAL which is 5% of the lipid composition. Assuming 50% of the DSPE-PEG-MAL faced inwards and if 100% of the peptide bound, then lipids had ~2.5% peptide (0.5uM). This improvement when attaching multiple copies of the inhibitory peptide to lipid was also seen with RI-OR2 [26]. Similarly, when Aβ peptide inhibitor RI-OR2-TAT was attached covalently to the surface of liposomes, to give a multivalent inhibitor, its potency as an aggregation inhibitor for Aβ was greatly improved [32]. When tested *in vivo*, Aβ oligomer and amyloid plaque levels were reduced in the cerebral cortex of 10-month-old APPswe/PS1ΔE9 mice following daily i.p. injections of RI-OR2-TAT over 21 days [31].

As anticipated, RI-AGO3 was resistant to proteolytic digestion. D-amino acids are typically resistant to proteolysis [26] and in RI-AG03, the backbone of the D-amino acids is reversed, resulting in an altered conformation of the trypsin cleavage site. Fluorescein-labelled RI-AG03 penetrated HEK cells, perhaps via cell mediated endocytosis or electrostatic processes. Polyarginine and other cell penetrating peptides can easily cross cell membranes without receptors [64] [65]. It is postulated that octa-arginine translocation is not limited to endocytosis but also through electrostatic and hydrophilic interactions by making hydrophilic holes in membranes [66] [67]. Direct penetration has been suggested whereby guanidinium side chains of Arginine nucleate transient pore formation [68]. Macropinocytosis may also be involved in the uptake of octa-Arginine, especially the larger poly-Arginine [69]. To escape endosomes, octa-Arginine may cause their membranes to leak, or utilise counteranion-mediated phase transfer [70]. Toxicity observed at high concentrations in **Figure 5b** may be associated with membrane leakage [71] [72].

75% of the genes implicated in human diseases have orthologues in *Drosophila*, this trait coupled with its genetic tractability and complex behavioural assays makes the fruit flies an attractive model for the study of neurodegenerative diseases and for screening drugs like RI-AG03. To test the efficacy of RI-AG03 in inhibiting Tau mediated phenotypes *in vivo*, a *Drosophila* model was employed. This model of tauopathy displays a characteristic “rough-eye” phenotype that is caused by a massive loss of underlying photoreceptor neurons induced by human-Tau (hTau) overexpression, hyperphosphorylation and aggregation [73] [39]. Importantly, this model has been used extensively to identify enhancers and suppressors of hTau-induced toxicity [74] [75] [76]. Our data suggests that RI-AG03 can suppress hTau mediated degeneration *in vivo* as is evident in the dose-dependent suppression of the “rough-eye” phenotype of treated Tau-expressing flies compared to age-matched untreated transgenics.

RI-AG03 similarly suppressed other hTau mediated phenotypes in *Drosophila*. Previous studies by co-authors have shown that the pan-neuronal expression of hTau decreases the lifespan of flies [38] and this was significantly improved by RI-AG03 treatment, at both low (0.08 μM) and high doses (0.8 μm) (**Figure 8**). Interestingly, this lifespan extension was not observed in the inhibitor-treated control lines, clearly due to the specificity of RI-AG03 in targeting monomeric and oligomeric human Tau proteins in the hTau-expressing transgenic flies. In addition, no decrease in lifespan was observed, demonstrating that the inhibitor is not toxic to Drosophila. As both the rough eye phenotype and shortened lifespan are believed to occur due to aggregation induced neurodegeneration of human Tau in these *Drosophila* models [39], this data collectively suggests that RI-AG03 was reducing Tau mediated aggregation *in vivo* in this model which manifested in the improved phenotypes.

### Conclusion

With the failure of Aβ targeting drugs in clinical trials, there is a renewed focus on Tau centric targets in the Alzheimer field. As aggregation is causally linked to most aspects of Tau toxicity, inhibition of Tau aggregation is likely to be an effective disease-modifying target. We therefore designed and developed a series of novel Tau aggregation inhibitors, of which, one RI-AG03 was found to be the most potent. It displays cell penetrability and is non-toxic both to cells in culture and *in vivo*. Importantly, treatment with this inhibitor potently suppresses Tau aggregation *in vitro* and significantly ameliorates Tau phenotypes *in vivo*. This collectively demonstrates the disease-modifying potential of inhibiting Tau aggregation and the utility of inhibitors such as RI-AG03 as therapeutic agents.

## METHODS

### Aggregation propensity

Aggrescan predicted aggregation propensity of experimental peptides by comparing the amino acid sequence to known hot spot sequences from 57 amyloidogenic proteins [77].

### Intrinsic solubility

Camsol Intrinsic predicted the intrinsic solubility profile of experimental peptides [45] [78] by evaluating amino acid hydrophobicity, aggregation propensity, charge and secondary structure propensity [79].

### Docking

Test peptides were docked onto Tau structures from the Protein Data Bank using Molsoft ICM-Pro (*version* 3.8-7). They did not include cell penetrating sequences due to applicability limitations in current protocols [80]. Water was kept tight and His, Pro, Asn, Gln and Cys side chains were optimised to correct orientation and H-bond network. Standard ICM-dock 3D grid potential maps used 0.5 Å grid spacing and docking probes were placed on the outer chains [81] [82]. Ligand atom types and charges were assigned by the Merck molecular force field. ‘Peptide docking mode’ forcefield was used to dock ligands onto Tau structures with maximum sampling effort factor [83] [84]. Flexible ring sampling (2) and biased probability Monte Carlo randomly selected independent subspaces based on predefined continuous probability distribution [85].

### Recombinant protein expression

Expression plasmid pRK-172 was transformed into *E. coli* BL21(DE3) cells and expressed as described [86]. Taniguchi-Watanabe received 4R1N in pRK-172 from Dr. Michel Goedert [87].

### Recombinant protein purification

Cells were resuspended in lysis buffer (50 mM PIPES, 1 mM EGTA, 1 mM DTT, 0.5 mM PMSF, 0.5 μg/ml leupeptin, pH 6.8) and lysed via sonication. Supernatant was supplemented with 1 % β-mercaptoethanol, boiled for 5 minutes and centrifuged at 27,000 × g. Supernatant was loaded onto a sulphopropyl (SP) sepharose column, pre-equilibrated in purification buffer (50 mM PIPES, 1 mM EGTA, 1 mM DTT, pH 6.8). TauΔ1-250 was eluted with purification buffer + 0.35 M NaCl. TauΔ1-250 was precipitated at 35% ammonium sulphate. Pellets were reconstituted in ddH_2_O and dialysed against Tris (30 mM), pH 7.5.

### Thioflavin-T fluorescence

Aggregation mix (Tau/peptide 20 μM, ThT 15 μM, dithiothreitol 1 mM, Tris 30 mM and heparin 5 μM pH 7.4) was incubated at 37°C with 10 seconds of shaking every 10 minutes for 24 hours. Fluorescence was measured (λex=442 nm, λem=483 nm) and data was normalised.

### TEM

10 μL samples were loaded onto Formvar/Carbon 300 mesh copper grids (Agar Scientific) for 3 minutes and then negatively stained with 10 μL of filtered 2% phosphotungstic acid, pH 2, for 2 minutes. Grids were examined using a JEOL JEM-1010 transmission electron microscope and were visualised and photographed at various magnifications.

### Circular dichroism

CD spectra were obtained using a Chirascan plus qCD spectrometer (Applied Photophysics) between 180 and 260 nm with a bandwidth of 1 nm and a path length of 2mm. Secondary structure estimation was conducted using the BeStSel web server.

### Peptide Liposomes

Peptide liposomes were prepared as described [31]. Excess unbound RI-AG03-C was removed by centrifugation at 171,000 g for 1 hour and the peptide-liposomes were resuspended in buffer. Peptide concentration was determined with a BSA assay and the amount of phospholipid was quantified using the Phospholipid C test from Fujifilm WAKO diagnostics USA.

### Statistics

Data are expressed as mean ± standard error of mean. ThT and *Drosophila* data was analysed using one factor repeated measures ANOVA with Tukey post hoc testing through IBM SPSS Statistics 23. For the Drosophila survival data, a Kaplan-Meier survival curve was plotted and a Log-rank (Mantel-Cox) test was performed on the data using GraphPad Prism [58].

### Enzyme stability

RI-AG03 (20 μM) was incubated at 37 °C for 24 hours with and without an equimolar concentration of Trypsin and run on SDS-PAGE gels.

### Cell culture

HEK-293 cell were maintained at 37 °C, 5% CO_2_ in Dulbecco’s Modified Eagle Medium/Ham’s F12 (DMEM/F12) at 1:1 supplemented with 10% FBS and 1% antibiotics (Streptomycin and Penicillin at 1:1).

### Cellular uptake

Sterile VWR pcs Cover Glass 13 mm were placed flat into wells of a 12 well plate. HEK-293 cells were seeded into individual wells at 20,000 cells, supplemented with 5(6)-carboxyfluorescein and incubated for 24 hours. Cells were fixed using 4% formaldehyde, washed with TBS and mounted to microscope slides using ProLong™ Gold Antifade Mountant. Samples were visualised on a Nikon Eclipse Ti fluorescent microscope using the FITC filter.

### Fruitfly Stocks

*Drosophila melanogaster* expressing either the retinal photoreceptor specific *GMR*-GAL4 driver or pan-neuronal driver *ElavC155*-GAL4, *UAS-*2N4RTau and Oregon-R (WT) flies were obtained from Bloomington Stock Centre, Indiana.

### *Drosophila*eye experiments

*UAS*-Tau flies were recombined to *GMR*-GAL4 driver to generate stable *GMR*-hTau lines (Bloomington Drosophila Stock Center 51361). *GMR*-GAL4 was used as the ‘healthy’ control. Progeny were reared on standard fly food supplemented with various RI-AG03 concentrations; 0.08μM, 0.8μM, 20μM and 40μM at 25°C. As controls, Tau and driver alone flies were incubated without the inhibitor. Freshly eclosed flies were monitored daily using a Nikon light microscope and 1-day old live flies were imaged (n=5/condition) and processed on ImageJ.

For SEM experiments, flies were euthanised, fixed in 2.5% glutaraldehyde in PBS and prepped for SEM with three 5-minute washes in PBS, before dehydrating the sample for 30 minutes at each ethanol concentration of 50%, 70%, 80%, 90%, 100%. Samples were then transferred into two changes of hexamethyldisilazane, after which hexamethyldisilazane was left to sublimate away. Samples were mounted onto JEOL SEM stubs under a stereo microscope before sputter coating with gold for 4 minutes on an Edwards 150A sputter coater. Samples were examined using a JEOL 5600 scanning electron microscope.

### *Drosophila*Survival Assays

*Elav*-GAL4 female flies were crossed with male flies overexpressing *UAS*-human 2N4R Tau (Full-length human Tau or hTau) or with gender-matched Oregon-R (WT) flies. Fly food was supplemented with either (0.08 μM or 0.8 μM RI-AG03). Ten cohorts of 10 male flies of each genotype were collected 0-5 days post-eclosion and transferred to new inhibitor-supplemented food twice a week. These were scored for deaths three times a week. A Kaplan-Meier survival curve was plotted and a Log-rank (Mantel-Cox) test was performed on the data using GraphPad Prism software [38].

## Acknowledgements

We thank the Sir John Fisher Foundation, Alzheimer’s Society and Alzheimer’s Research UK for financial support of this research.

**Correspondence and requests for materials should be addressed to AM.**

## SUPPLEMENTARY

**Figure S1:**
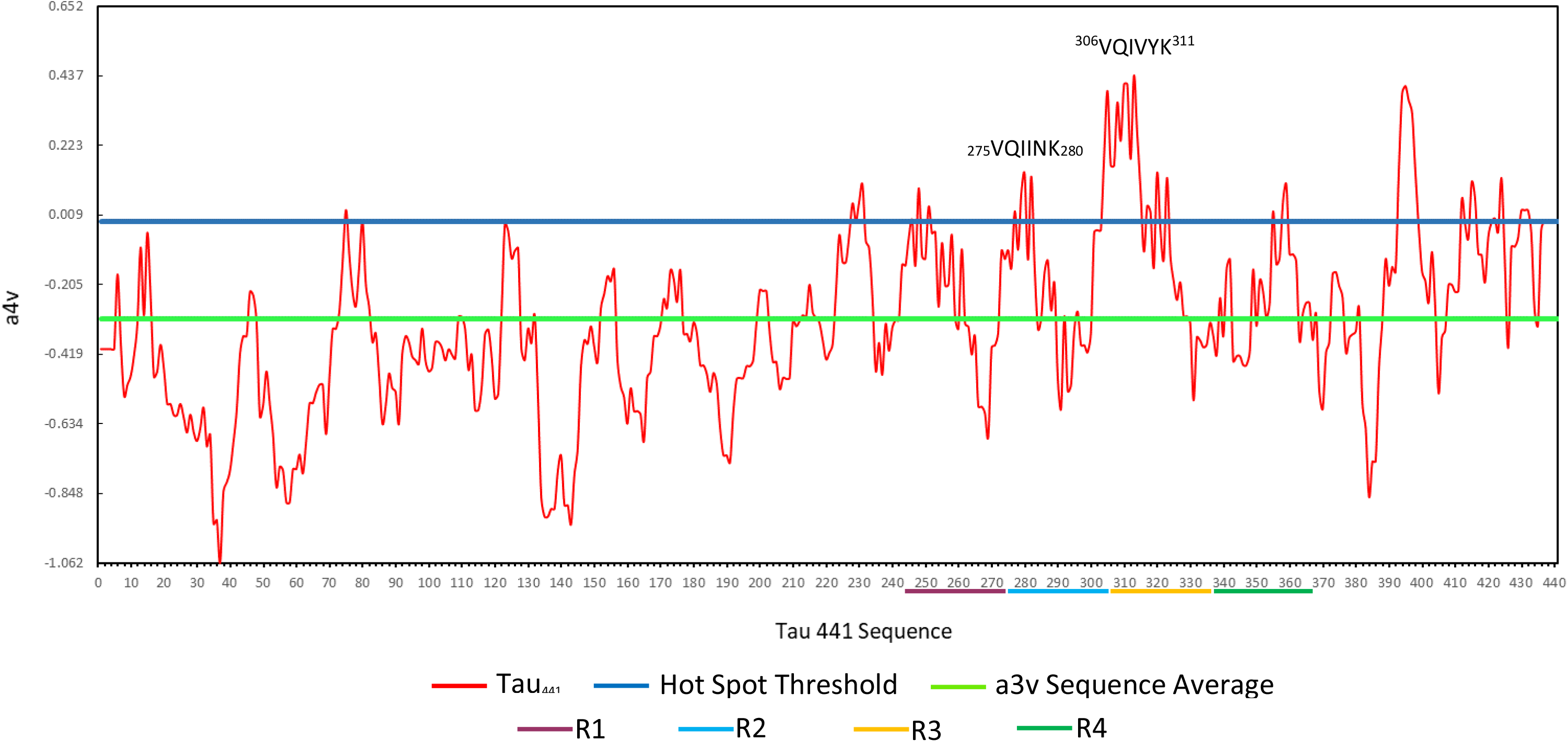
AGGRESCAEN average aggregation propensity for Tau_441_ residues. VQIVYK exceeds the hot spot threshold (blue line) significantly. R1-4 signifies the repeat regions in Tau. Hexapeptides exceeding the hot spot threshold are highlighted as residues likely to aggregate.

**Figure S2:**
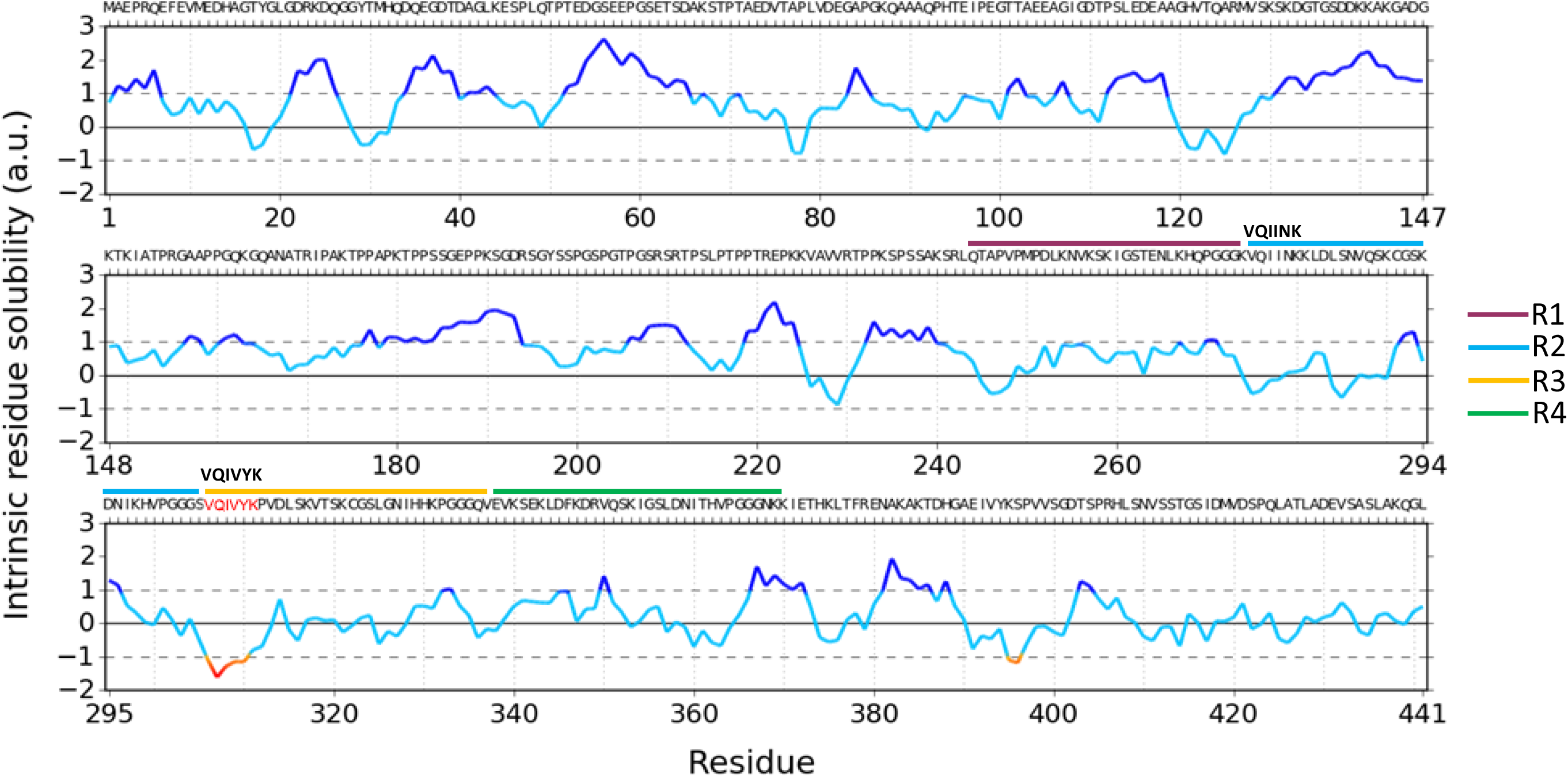
CamSol Intrinsic residue solubility for Tau_441_ residues. VQIVYK contains hydrophobic residues valine and isoleucine. The hexapeptide is identified with a red line, suggesting an aggregation hot spot. Blue lines suggest soluble regions. R1-4 signifies the repeat regions in Tau.

**Table S3:**
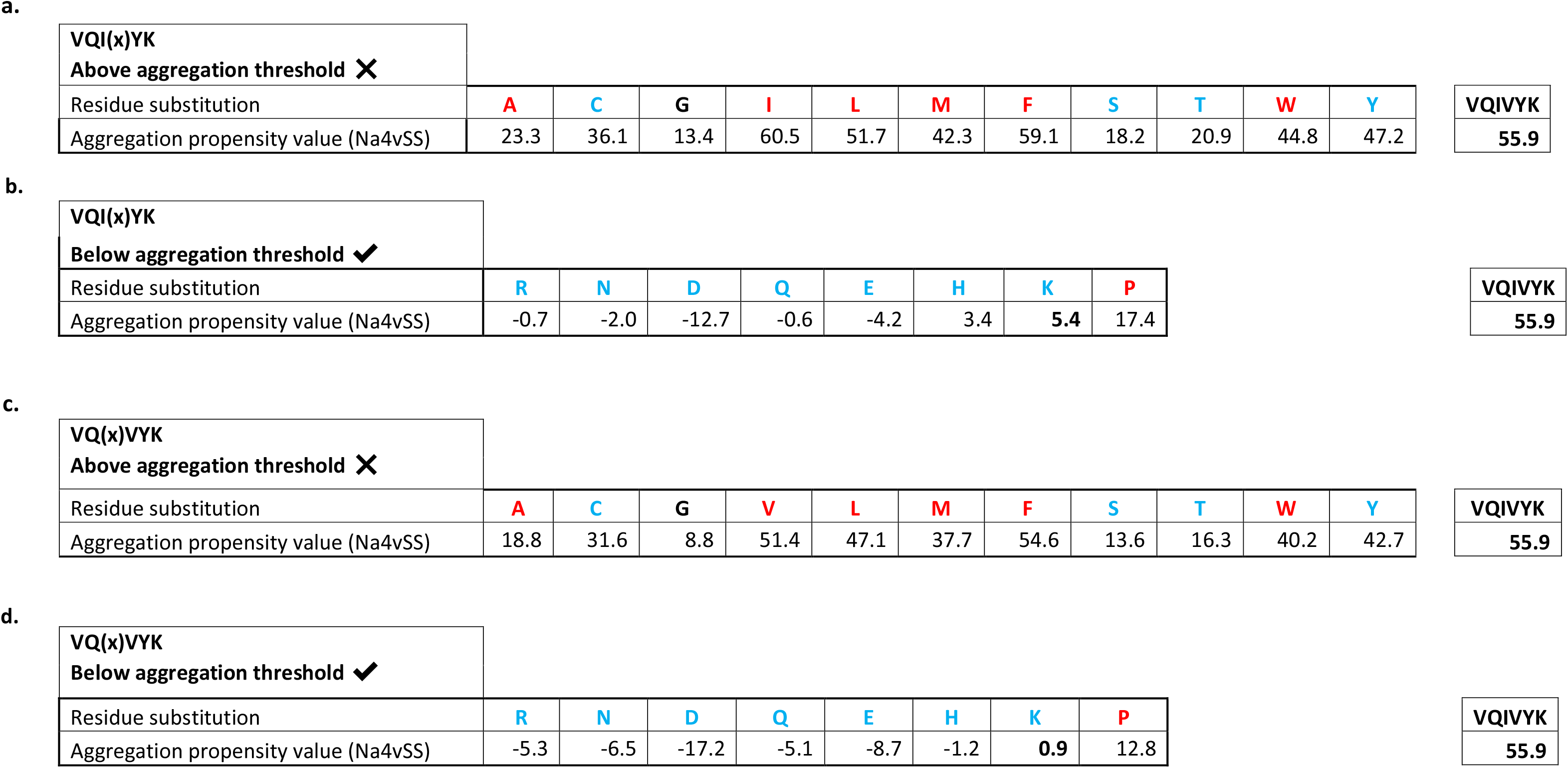
Summarises the average aggregation propensity values for VQI(x)YK and VQ(x)VYK sequences, using replacements at (x) with different amino acids for comparison. **a+c:** The conformations which were predicted to aggregate, **b+d:** The sequences which were predicted not to aggregate. Replacement at x with R, N, D, Q, E, H, K and P massively reduced the average aggregation propensity. Residues coloured in red, blue and black are hydrophobic, hydrophilic or neutral, respectively.

|  |  |  |  |  |  |  |  |  |  |  |  |  |
| --- | --- | --- | --- | --- | --- | --- | --- | --- | --- | --- | --- | --- |
| VQI(x)YK<br>Above aggregation threshold ✕ |  |  |  |  |  |  |  |  |  |  |  | VQIVYK<br>55.9 |
| Residue substitution | A | C | G | I | L | M | F | S | T | W | Y |  |
| Aggregation propensity value (Na4vSS) | 23.3 | 36.1 | 13.4 | 60.5 | 51.7 | 42.3 | 59.1 | 18.2 | 20.9 | 44.8 | 47.2 |  |

b.
|  |  |  |  |  |  |  |  |  |  |
| --- | --- | --- | --- | --- | --- | --- | --- | --- | --- |
| VQI(x)YK |  |  |  |  |  |  |  |  | VQIVYK |
| Below aggregation threshold ✓ |  |  |  |  |  |  |  |  |  |
| Residue substitution | R | N | D | Q | E | H | K | P |  |
| Aggregation propensity value (Na4vSS) | -0.7 | -2.0 | -12.7 | -0.6 | -4.2 | 3.4 | 5.4 | 17.4 | 55.9 |
| c. |  |  |  |  |  |  |  |  |  |  |  |  |
| VQ(x)VYK<br>Above aggregation threshold ✕ |  |  |  |  |  |  |  |  |  |  |  | VQIVYK<br>55.9 |
| Residue substitution | A | C | G | V | L | M | F | S | T | W | Y |  |
| Aggregation propensity value (Na4vSS) | 18.8 | 31.6 | 8.8 | 51.4 | 47.1 | 37.7 | 54.6 | 13.6 | 16.3 | 40.2 | 42.7 |  |

d.
|  |  |  |  |  |  |  |  |  |  |
| --- | --- | --- | --- | --- | --- | --- | --- | --- | --- |
| VQ(x)VYK<br>Below aggregation threshold ✓ |  |  |  |  |  |  |  |  | VQIVYK<br>55.9 |
| Residue substitution | R | N | D | Q | E | H | K | P |  |
| Aggregation propensity value (Na4vSS) | -5.3 | -6.5 | -17.2 | -5.1 | -8.7 | -1.2 | 0.9 | 12.8 |  |

**Table S4:**
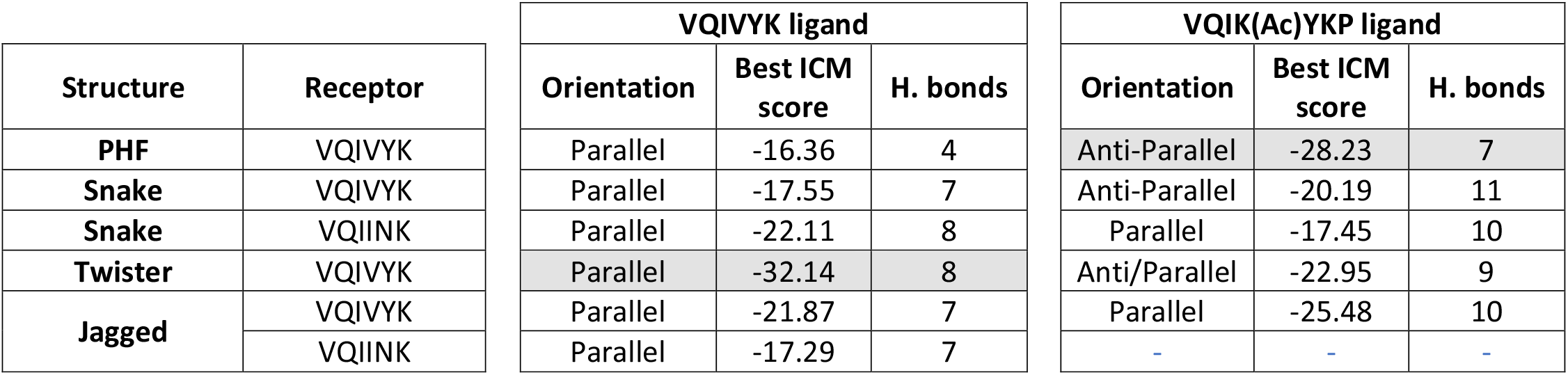
Summarises the strongest energy, orientation and hydrogen bond count for VQIVYK and VQIK(Ac)YKP docked to PDB: 5o3l PHF, 6QJH snake filament, 6QJM twister filament and 6QJP jagged filament. The strongest energy binds are highlighted in green.

**Table S5:**
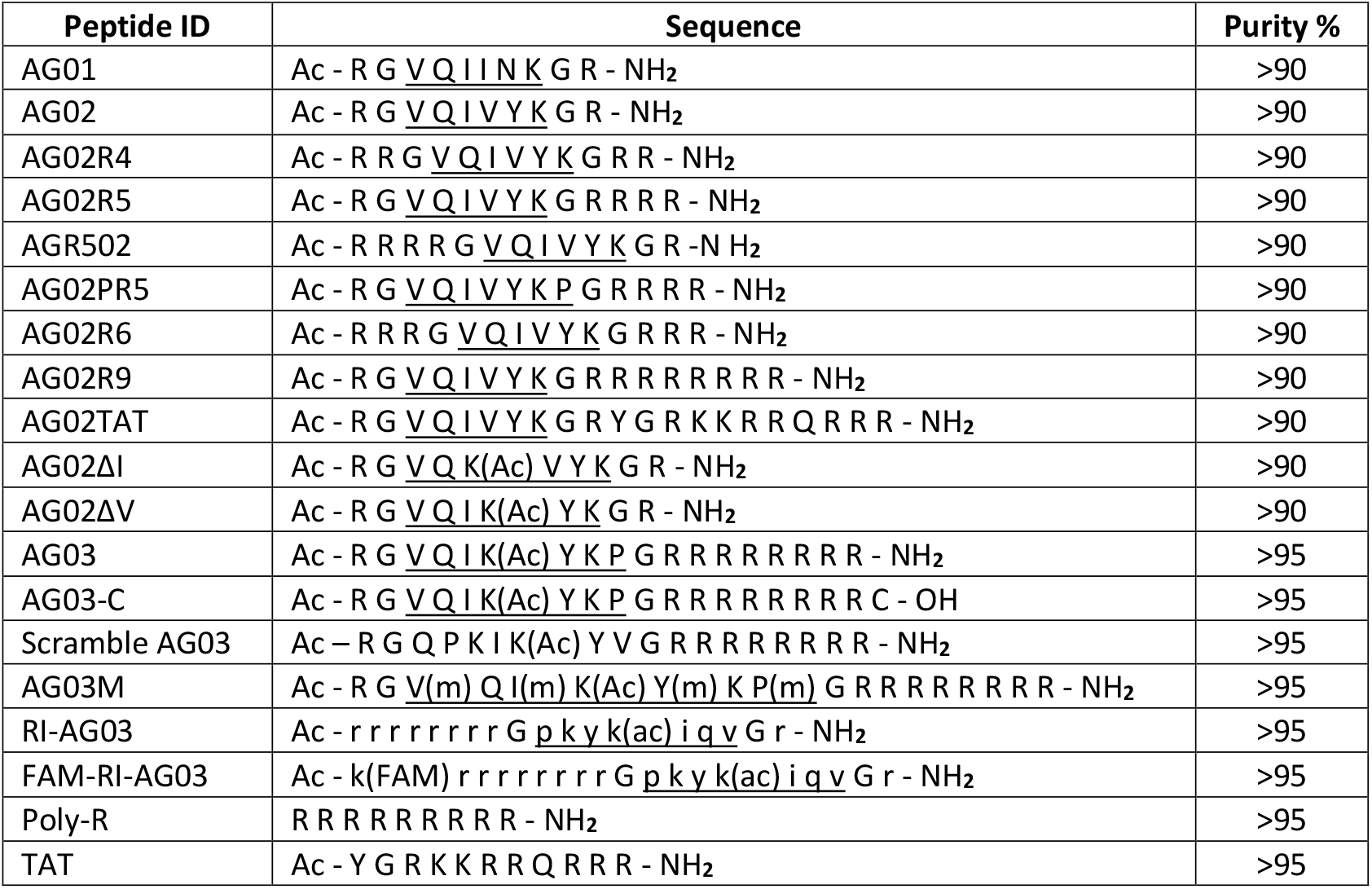
These are the experimental peptide inhibitor designs used in this research. Underlined portions indicate the predicted Tau binding regions. Peptides were synthesised by Peptide Synthetics (Fareham, UK) and Severn Biotech (Kidderminster, UK). Purity was determined by HPLC-MS. Flanking the peptide binding sequence with RG and GR spacers may improve binding ability by distancing other amino acids, e.g. cell penetrating sequences which could potentially interfere. Activity was enhanced by acetylating and amidating the peptide terminals. Stability of AG03 was enhanced with either N-methylation or retro-inversion.

**Figure S6:**
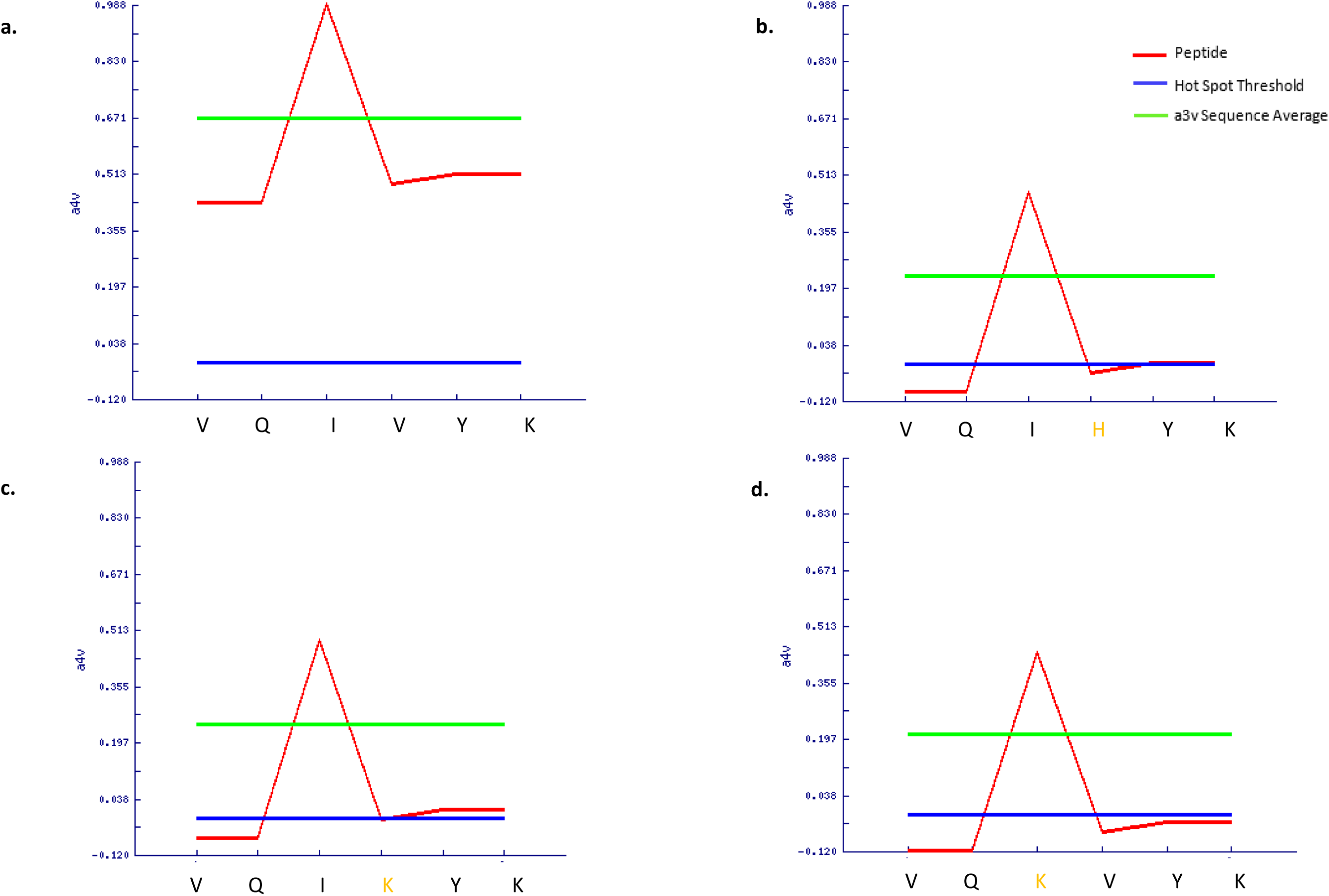
AGGRESCAN average aggregation propensity for Tau residues 301-315 with a replaced amino acid indicated in orange. **a:** Native VQIVYK; **b:** VQI**H**YK; **c:** VQI**K**YK; **d:** VQ**K**VYK. Replacing isoleucine or valine greatly reduces the aggregation propensity.

**Figure S7:**
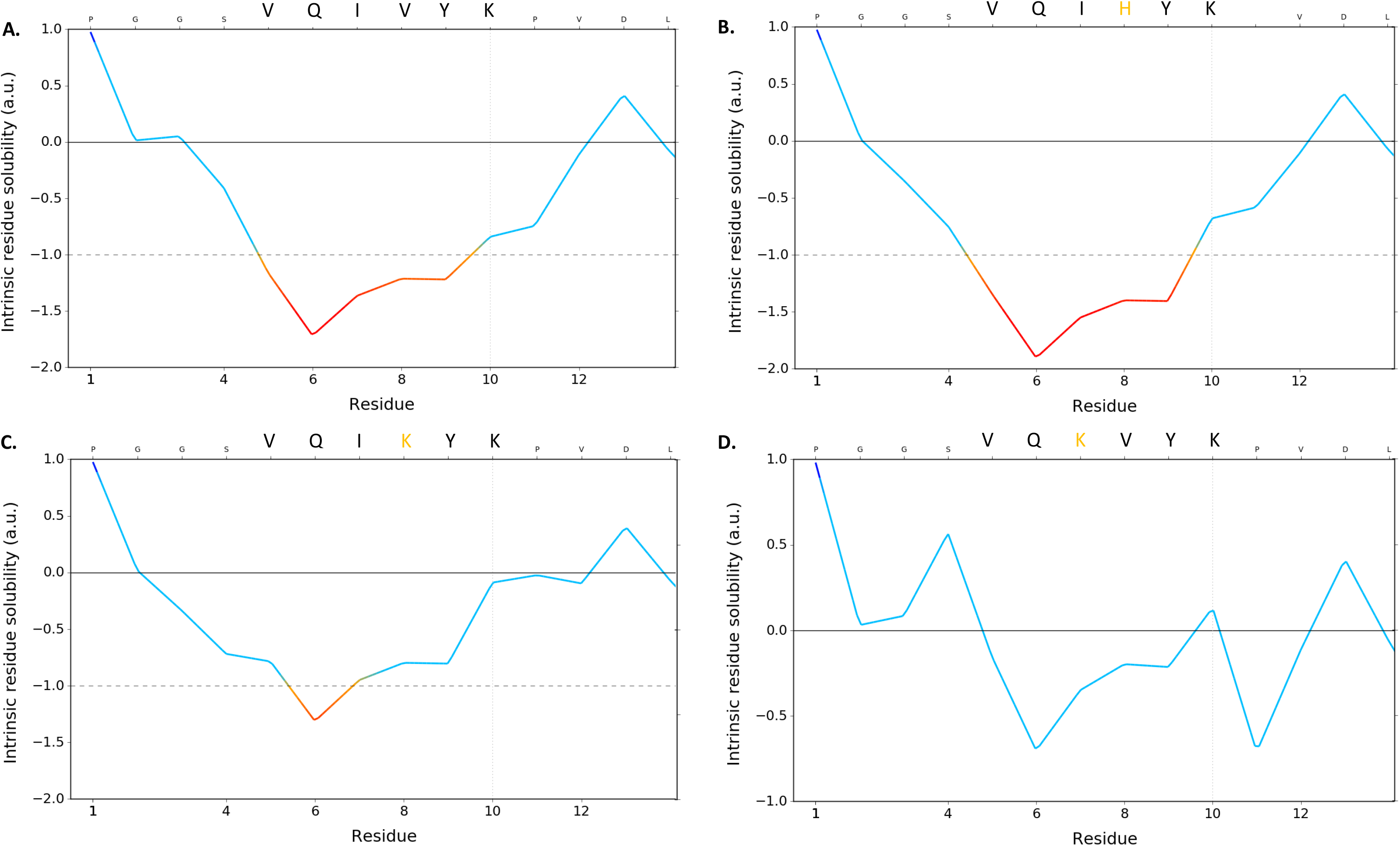
CamSol Intrinsic residue solubility for Tau residues 301-315 with a replaced amino acid indicated in orange. **A:** Native VQIVYK; **B:** VQI**H**YK; **C:** VQI**K**YK; **D:** VQ**K**VYK. Isoleucine appears to be more important for influencing intrinsic solubility than valine is in the VQIVYK sequence.

**Figure S8:**
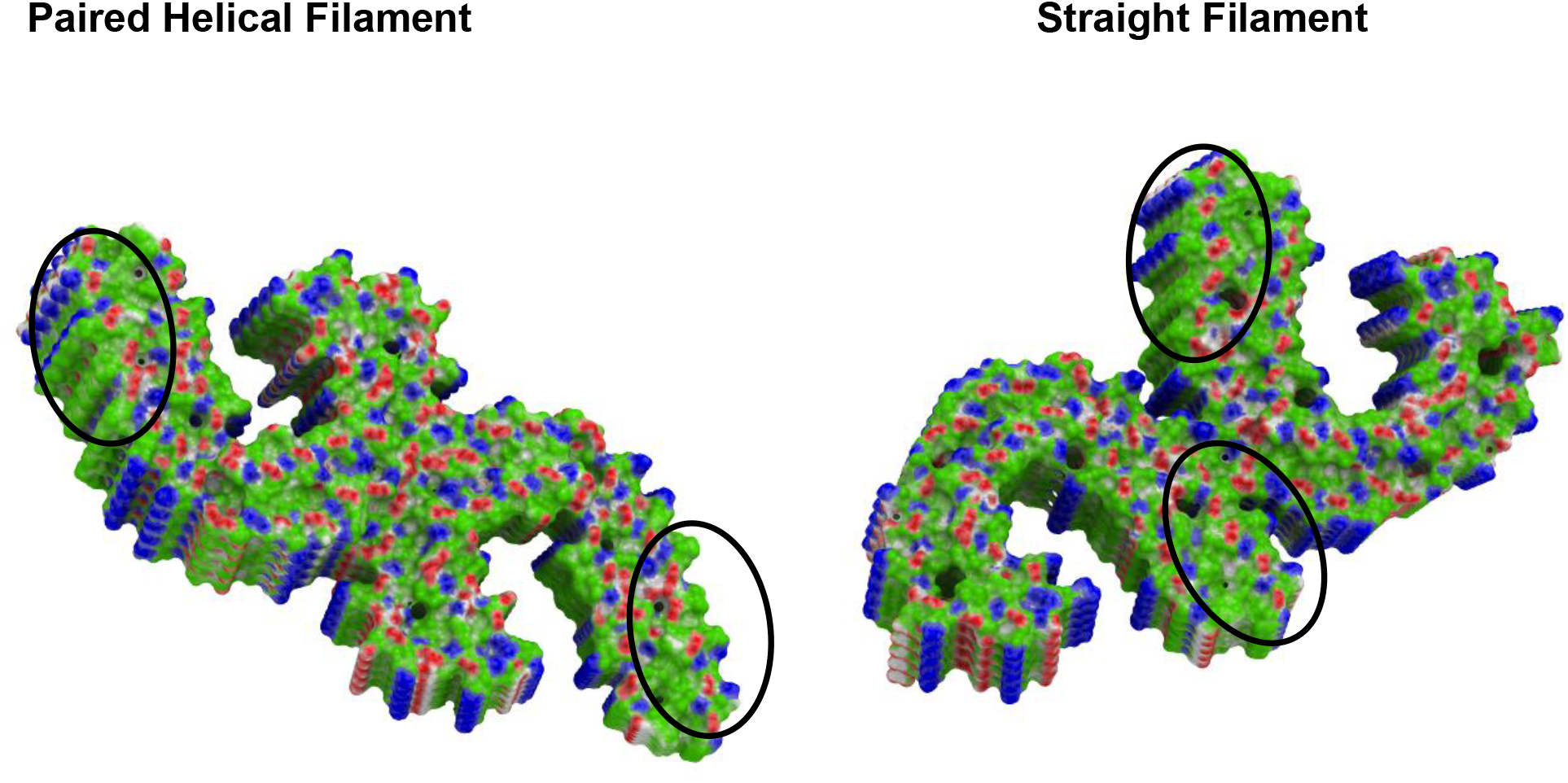
Demonstrates the binding properties of PDB 5o3l PHF and PDB 5o3t SF. This includes hydrophobic areas (green), hydrogen bond acceptor potential (red) and hydrogen bond donor potential (blue). The VQIVYK region is circled in black and mainly contains hydrogen bonding donors on the lateral, whereas perpendicular to the fibril axis are mainly hydrogen bonding acceptors.

**Figure S9:**
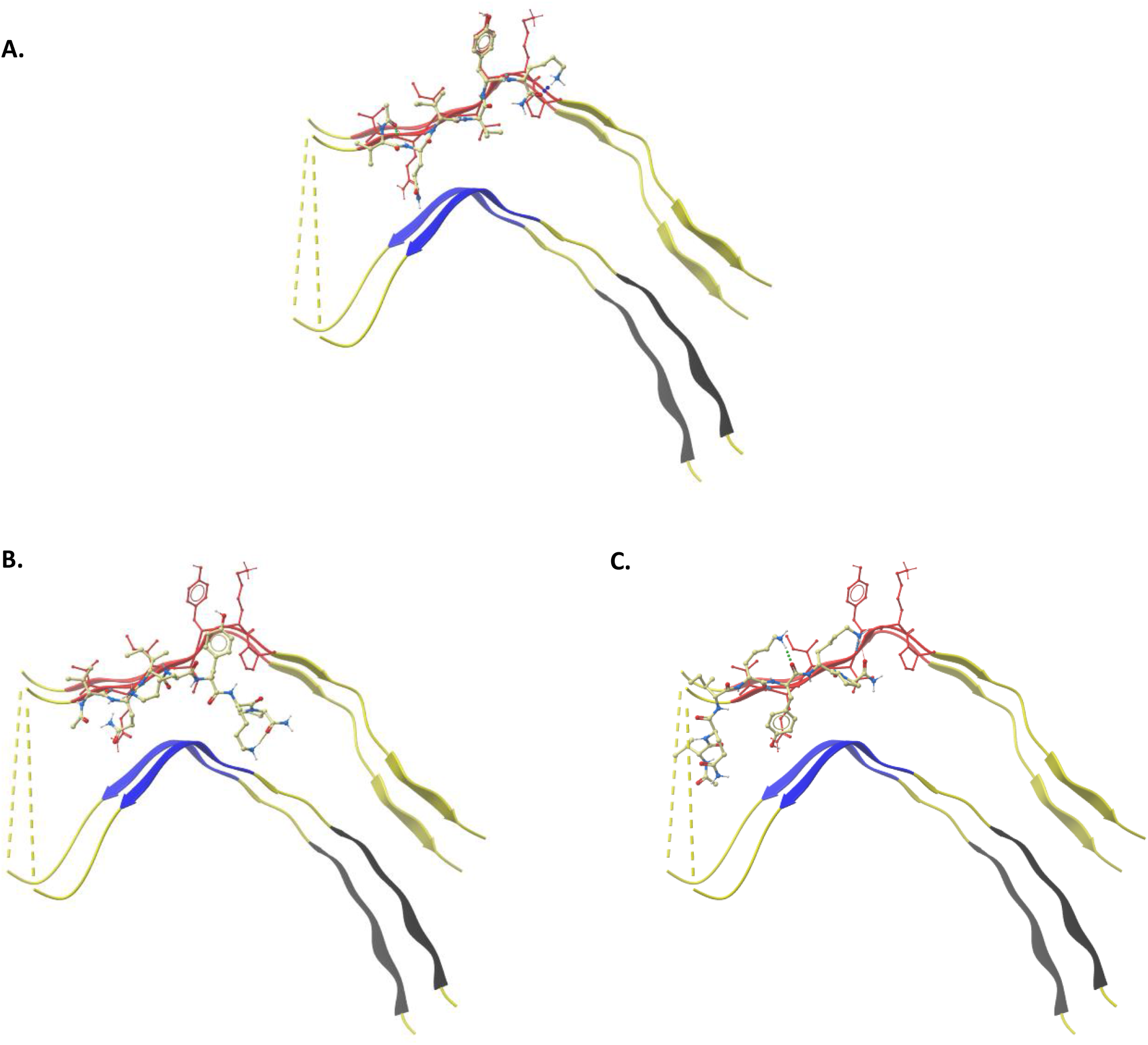
Docked peptides to PDB 6QJM heparin-induced 2N4R Tau twister filament; each binding to the ^306^VQIVYK^311^ sequence. **A:** VQIVYK binding in parallel to fibril, **B:** VQIK(Ac)YKP binding in paralell to the fibril but slightly shifted, **C:** VQIK(Ac)YKP binding anti-parallel to fibril.

**Figure S10:**
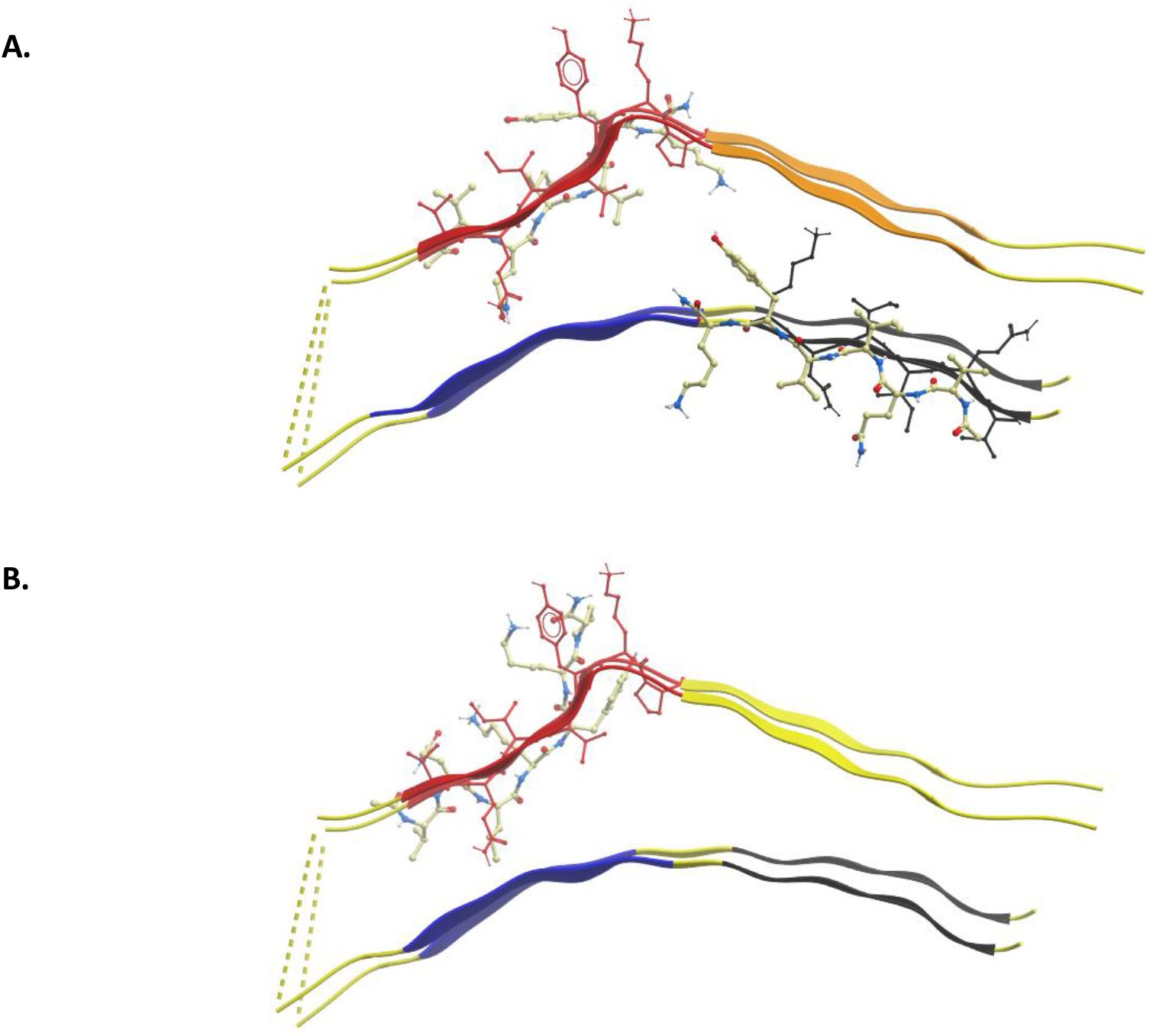
Docked peptides to PDB 6QJP heparin-induced 2N4R Tau jagged filament. **A:** VQIVYK binding in parallel to fibril at the ^306^VQIVYK^311^ and ^275^VQIINK^280^ positions, **B:** VQIK(Ac)YKP binding in parallel at the VQIVYK position.

**Figure S11:**
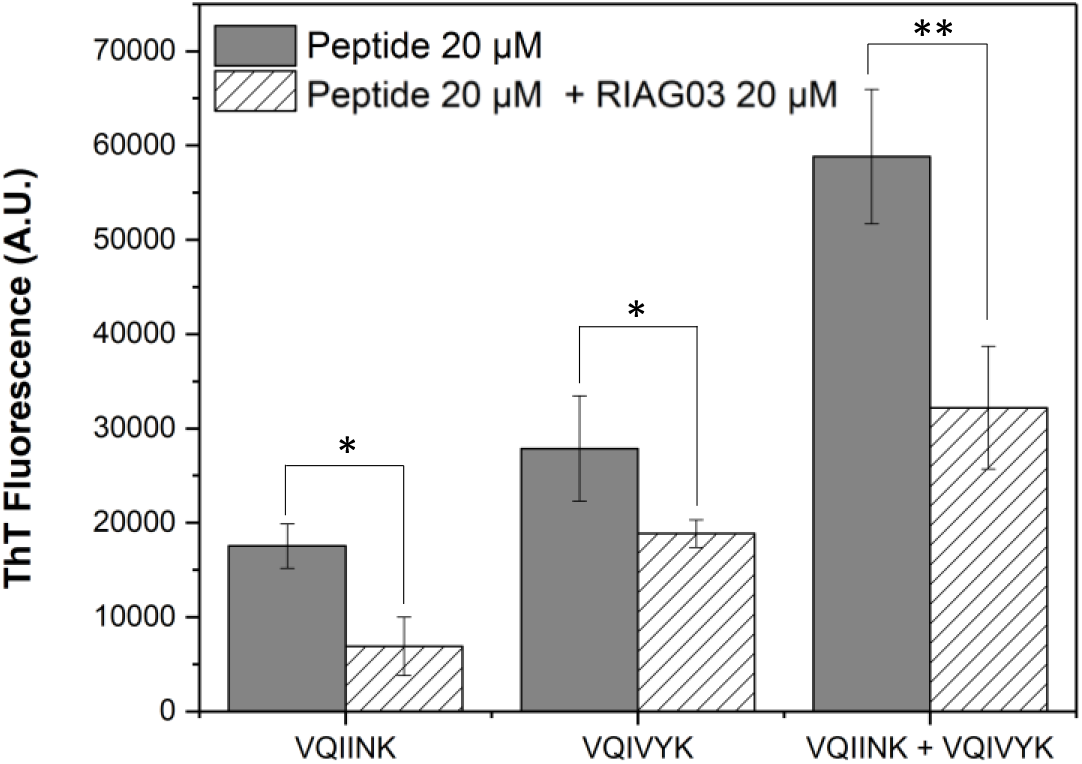
Thioflavin data using a Synergy2 plate reader. **a:** End-point (24 hr) aggregation of VQIINK (20 μM), VQIVYK (20 μM) and a combination of both with and without RI-AG03 [rrrrrrrrg-pkyk(Ac)iqv-gr] (20 μM). P > 0.05, * P ≤ 0.05, ** P ≤ 0.01, *** P ≤ 0.001.

